# Repulsive interactions instruct synaptic partner matching in an olfactory circuit

**DOI:** 10.1101/2025.03.01.640985

**Authors:** Zhuoran Li, Cheng Lyu, Chuanyun Xu, Ying Hu, David J. Luginbuhl, Asaf B. Caspi-Lebovic, Jessica M. Priest, Engin Özkan, Liqun Luo

## Abstract

Neurons exhibit extraordinary precision in selecting synaptic partners. Although cell-surface proteins (CSPs) mediating attractive interactions between developing axons and dendrites have been shown to instruct synaptic partner matching^1,2^, the degree to which repulsive interactions play a role is less clear. Here, using a genetic screen guided by single-cell transcriptomes^3,4^, we identified three CSP pairs—Toll2–Ptp10D, Fili–Kek1, and Hbs/Sns– Kirre—in mediating repulsive interactions between non-partner olfactory receptor neuron (ORN) axons and projection neuron (PN) dendrites in the developing *Drosophila* olfactory circuit. Each CSP pair exhibits inverse expression patterns in the select ORN-PN partners. Loss of each CSP in ORNs led to similar synaptic partner matching deficits as the loss of its partner CSP in PNs, and mistargeting phenotypes caused by overexpressing one CSP could be suppressed by loss of its partner CSP. All CSP pairs are also differentially expressed in other brain regions. Together, our data reveal that multiple repulsive CSP pairs work together to ensure precise synaptic partner matching during development by preventing neurons from forming connections with non-cognate partners.

## Main

A fundamental question in neural development is how the vast number of neurons precisely select their synaptic partners to form functional circuits. Neural circuit wiring involves multiple coordinated developmental steps: axon guidance to target regions, dendrite patterning, and synaptic partner matching followed by synaptogenesis^5–7^. Even though axon guidance and dendrite patterning can greatly reduce the number of potential partners a neuron encounters at a given time and region^8^, a developing axon must select specific partners among multiple nearby non-partners^1,2^. The mechanisms by which neural systems reduce multiple candidate synaptic partners to a specific one remain poorly understood.

It is well established that axon guidance involves both attraction towards the target region and repulsion away from non-target regions^9,10^. Repulsion mediated by cell-surface proteins (CSPs) is also used in establishing topographic maps^11^, subregion target selection^12^, and dendritic and axonal self-avoidance^2^. However, most known CSPs that instruct the final steps of synaptic partner selection act through attraction. These include homophilic attraction of Teneurins (Ten-m and Ten-a) in *Drosophila* olfactory and neuromuscular systems^13,14^, heterophilic attractions among members of the immunoglobulin superfamily of CSPs in multiple *Drosophila* circuits^15–22^, and homophilic attraction mediated by immunoglobulin^23^ or cadherin^24,25^ families of CSPs in the vertebrate retina. The few repulsion examples include *Drosophila* motor axon target selection controlled by Wnt4 from non-target muscles^26^ and olfactory neuron target selection by Fish-lips (Fili) from non-cognate partners^27^. How general repulsion is employed as a guiding force in synaptic partner matching remains to be examined.

In the *Drosophila* olfactory circuit, axons of about 50 types of olfactory receptor neurons (ORNs) form one-to-one precise synaptic connections with dendrites of 50 types of projection neurons (PNs) in 50 glomeruli in the antennal lobe^28^. During development, PN dendrites coarsely pattern the antennal lobe first^29,30^. While extending across the antennal lobe prepatterned by PN dendrites, each ORN axon sends multiple transient branches along its trajectory. ORN axon branches that contact partner PN dendrites are stabilized and branch further, whereas the rest retract^31,32^. Since synaptic partner matching involves retraction of transient ORN axon branches in contact with non-partner PNs, we aimed to identify repulsive CSPs that might function to prevent the formation of misconnections between non-partner PNs and ORNs.

### Inverse expression of three CSP pairs

VA1d and VA1v are neighboring glomeruli that sense distinct pheromones^33,34^. Known homophilic attraction molecules mediating matching between synaptic partners, Ten-m and Ten-a, cannot distinguish VA1d-PNs and VA1d-ORNs from VA1v-PNs and VA1v-ORNs, as they all express Ten-m at high levels and Ten-a at low levels^13^. We hypothesized that additional CSPs are differentially expressed and instruct synaptic partner matching in these adjacent glomeruli. To identify such CSPs, we performed a genetic screen focusing on PN-ORN matching in the VA1d and VA1v glomeruli (Fig. 1a). We first analyzed the existing single-cell transcriptome data for developing PNs and ORNs^3,4^ at 24–30h after puparium formation (APF), shortly before matching between ORN axons and PN dendrites occurs. We focused on CSPs (including both transmembrane and secreted proteins^35^) that are differentially expressed in VA1d-PNs and VA1v-PNs, or VA1d-ORNs and VA1v-ORNs. We narrowed down to 36 candidate genes (see below) with assistance from existing literature, including the list of top 100 CSPs enriched in developing antennal lobes revealed by proteomic profiling^36^. We then performed tissue-specific RNA interference (RNAi) against candidate genes selectively in PNs, ORNs, and/or all neurons.

**Fig. 1.**
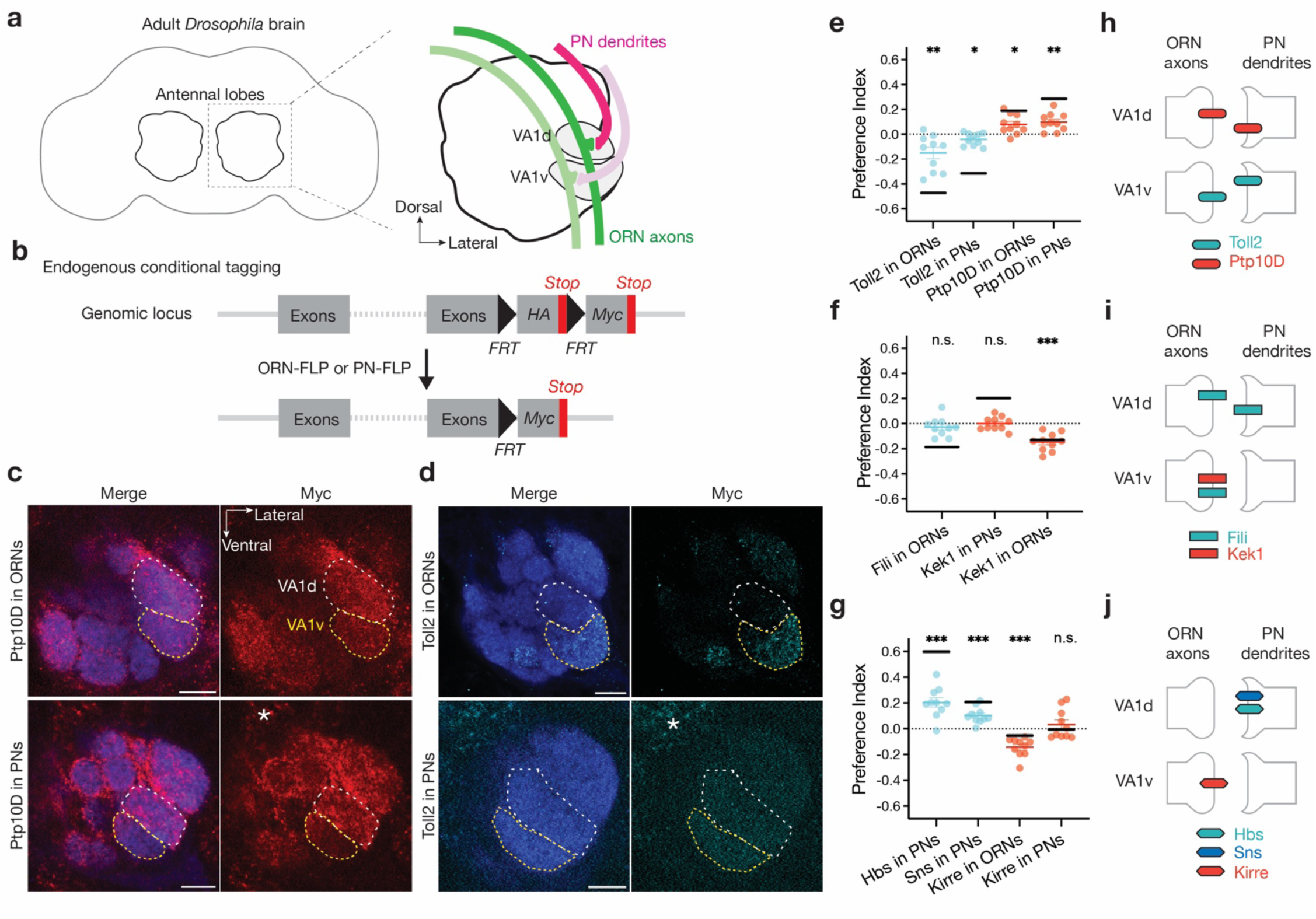
Inverse expression of three CSP pairs in the VA1d and VA1v glomeruli. **a,** Adult *Drosophila* brain and the antennal lobe schematics. Axons of VA1d-ORNs and VA1v-ORNs (green) match with dendrites of VA1d-PNs and VA1v-PNs (magenta), respectively. **b,** Schematic of conditional tagging of CSPs to reveal their endogenous protein expression pattern in whole brain (top) before—and in specific cell types (bottom) after—FLP-mediated recombination. **c, d,** Confocal images showing neuropil staining by N-cadherin antibody (blue) and Myc staining of tagged endogenous Ptp10D (red in **c**) and Toll2 (cyan in **d**) using ORN-specific FLP (top tow) or PN-specific FLP (bottom row). The VA1d and VA1v glomeruli are outlined in white and yellow, respectively, based on N-cadherin staining. Example images of other proteins are shown in Extended Data Fig. 2c. **e–g,** Quantification of the preference index (Methods) of RNA (black horizontal lines) and protein (colored data points: red for Ptp10D, Kek1, and Kirre; cyan for Toll2, Fili, Hbs, and Sns) expression levels in ORNs and PNs that innervate the VA1d versus VA1v glomerulus. Preference index > 0 means that expression level in VA1d is higher than in VA1v. Preference index for proteins was calculated based on Myc staining intensity throughout the glomeruli at 42–48h APF (data in **c, d,** Extended Data Fig. 2c). For all genotypes, n ≥ 10. Cyan or red lines indicate geometric mean. One-sample t-test comparing to zero. Preference index for mRNAs was calculated based on average expression levels in VA1d-ORN/PNs versus VA1v-ORN/PNs based on the single-cell-transcriptome data at 24–30h APF^3,4^ (black lines). **h–j,** Summary of relative CSP expression levels in the VA1d and VA1v PN-ORN pairs during development based on mRNA and protein data in **e–g** (see Extended Data Fig. 2 for caveats of each). If the expression level in VA1d-and VA1v-ORNs/PNs is significantly different, we only drew the bars in the cell types with higher expression level as a simplification. In this and all subsequent figures, * P < 0.05; ** P < 0.01; *** P < 0.001; n.s. not significant, Scale bars = 10 µm unless otherwise noted.

In wild-type animals, axons of VA1d-ORNs and VA1v-ORNs only innervated the VA1d and VA1v glomeruli, respectively (Extended Data Fig. 1a, d). 14 of the 36 candidate genes from the screen showed mistargeting phenotypes in either VA1d-ORNs or VA1v-ORNs (Extended Data Fig. 1; Extended Data Table 1; companion manuscript^37^). We note that loss-of-function of single CSP usually resulted in subtle phenotypes: mistargeting of a small fraction of axons or dendrites. In the companion manuscript, we showed that only by simultaneously manipulating multiple CSPs in ORNs could we substantially change their matching specificity^37^. Among these candidate genes, three pairs of CSPs—Toll2–Ptp10D, Fili–Kek1, and Hbs/Sns–Kirre—exhibited largely inverse expression patterns in ORN-PN synaptic partners based on single-cell transcriptome data, particularly in ORNs and PNs that target the VA1d and VA1v glomeruli (Extended Data Fig. 2a). For example, *Toll2* is more highly expressed in VA1v-ORNs than VA1d-ORNs, whereas its partner *Ptp10D* is more highly expressed in VA1d-PNs than VA1v-PNs (Extended Data Fig. 2a, left column). Such inverse expression patterns suggest a potential role for these CSP pairs to promote repulsion during synaptic partner matching. We thus focused the rest of our study on these three CSP pairs.

To validate the mRNA-based inverse expression patterns (Extended Data Fig. 2a), we examined the endogenous protein expression levels at 42–48h APF, when glomerular identities first become identifiable (and the matching between partner ORN axons and PN dendrites is mostly complete). To determine cell-type-specific expression patterns, we knocked into the endogenous loci of the three CSP pairs a modified conditional tag^38,36^ (Fig. 1b). In the absence of the FLP recombinase, these proteins were tagged with HA and no Myc signal was detected in the antennal lobe (Extended Data Fig. 2b). With ORN-specific FLP or PN-specific FLP, we could visualize endogenous protein expression only in ORNs or PNs by Myc staining (Fig. 1c, d, Extended Data Fig. 2c). We found that Toll2 (also named 18-wheeler) and Ptp10D (protein tyrosine phosphatase 10D) were differentially expressed across the antennal lobe (Supplementary Videos 1–7). In the VA1d and VA1v glomeruli, Toll2 exhibited higher expression in VA1v-PNs and VA1v-ORNs, whereas Ptp10D had higher expression in VA1d-PNs and VA1d-ORNs (Fig. 1c, e). Using similar conditional tags, we found that Kek1 (Kekkon 1) exhibited higher expression in VA1v-ORNs than in VA1d-ORNs and low expression in both VA1d-PNs and VA1v-PNs (Fig. 1f and Extended Data Fig. 2c). Fili did not show preferential expression in VA1d-ORNs and VA1v-ORNs but exhibits higher expression in VA1d-PNs than VA1v-PNs based on data from a previous study^27^. For the third CSP pair, Hbs (Hibris) and Sns (Sticks and stones) were minimally expressed in ORNs and exhibited higher expression in VA1d-PNs than VA1v-PNs, whereas Kirre (Kin of irre) exhibited higher expression in VA1v-ORNs than VA1d-ORNs (Fig. 1g and Extended Data Fig. 2c). Using the same preference index to quantify the relative expression in ORNs or PNs that target VA1d vs. VA1v glomerulus, we found that mRNA and protein expression patterns were mostly consistent for all three CSP pairs (Fig. 1e–g). We note that the magnitudes of differential expression for some CSPs, while significant, were modest; nevertheless, our genetic analysis below suggests that such differential expression was indeed used to instruct synaptic partner matching.

In summary, based on the relative expression levels of mRNAs and proteins, the Toll2–Ptp10D, Fili–Kek1, and Hbs/Sns–Kirre pairs are expressed in inverse patterns in PN-ORN partners at the VA1d and/or VA1v glomeruli (Fig. 1h–j). Furthermore, all these CSPs are present at the terminals of ORN axons and/or PN dendrites at the nascent glomeruli, consistent with their playing a role in synaptic partner matching.

### Loss of Toll2 or Ptp10D disrupts partner matching

We first examined the function of Toll2 and Ptp10D in PN-ORN synaptic partner matching. Ptp10D is an evolutionarily conserved member of the Type III receptor tyrosine phosphatase (RPTP) family (Fig. 2a) involved in axon guidance at the midline, tracheal tube formation, and cell competition, and was reported to be a receptor for the CSP Sas (Stranded at second)^39–43^. However, single-cell transcriptomic data indicate that Sas is minimally expressed in the antennal lobe^3,4^, suggesting the existence of additional Ptp10D-interacting CSPs.

**Fig. 2.**
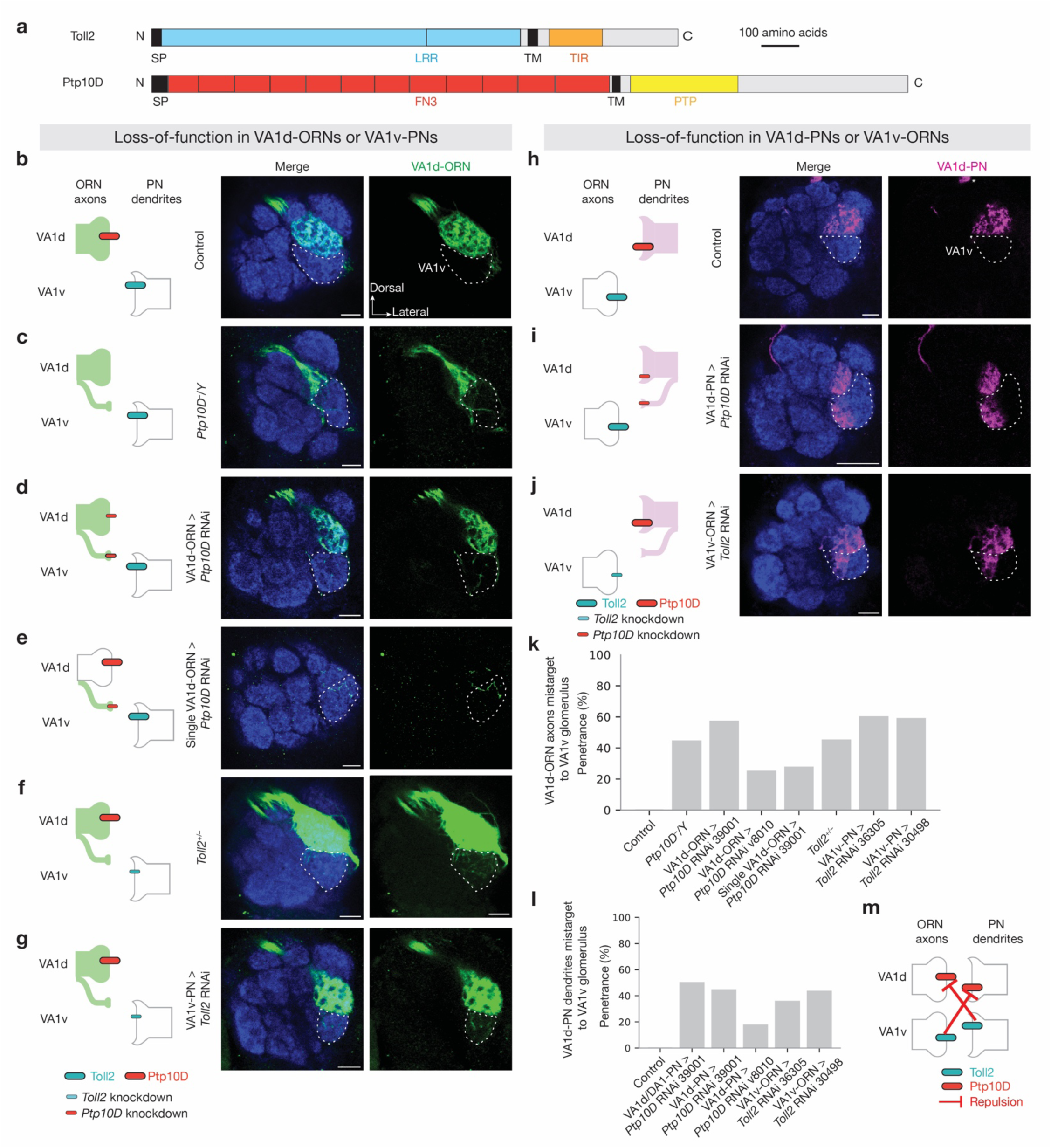
Loss of Ptp10D or Toll2 causes similar mismatching between non-partner PNs and ORNs. **a,** Domain composition of *Drosophila* Toll2 and Ptp10D. TM, transmembrane domain; N, N-terminus; C, C-terminus; SP, signal peptide; LRR, leucine-rich repeat; TIR, Toll/interleukin-1 receptor; FN3, fibronectin type III; PTP, protein tyrosine phosphatase domain. **b–j,** Left column shows experimental schematic. Middle and right columns are confocal images of adult antennal lobes showing neuropil staining by N-cadherin antibody (blue) and VA1d-ORN axons (green in **b–g**) or VA1d-PN dendrites (magenta in **h–j**). The VA1v glomerulus is outlined based on N-cadherin staining. Control VA1d-ORN axons innervate the VA1d glomerulus (**b**). Some VA1d-ORN axons mistarget to the VA1v glomerulus in *Ptp10D* hemizygous mutant (**c**), *Ptp10D* RNAi expressed in all (**d**) or single (**e**) VA1d-ORNs, *Toll2* heterozygous mutant (**f**), or *Toll2* RNAi expressed in VA1v-PNs (**g**). Control VA1d-PN dendrites only innervate the VA1d glomerulus (**h**). Some VA1d-PN dendrites mistarget to the VA1v glomerulus when *Ptp10D* RNAi was expressed in VA1d-PNs (**i**) or when *Toll2* RNAi was expressed in VA1v-ORNs (**j**). * designates PN cell bodies. **k, l,** Penetrance of the mistargeting phenotypes in **b–g** (**k**) and **h–j** (**l**). Mistargeting is counted if at least one trackable axon or dendrite innervate inside the VA1v glomerulus from the 3D confocal stacks. For all genotypes, n ≥ 10. **m,** Schematic summary for the function of Ptp10D and Toll2 in VA1d and VA1v glomeruli. **—|**, repulsive signaling from the open end (sender) to the end with a perpendicular line (receiver).

To validate the *Ptp10D* RNAi phenotypes from our screen (Extended Data Fig. 1b), we labeled VA1d-ORN axons in *Ptp10D* hemizygous mutant animals and observed similar phenotype as pan-ORN *Ptp10D* RNAi: VA1d-ORN axons mistargeted to the VA1v glomerulus (Fig. 2b, c, k). Given the high Ptp10D expression in VA1d-ORNs (Fig. 1h), we tested whether Ptp10D is autonomously required in VA1d-ORNs for their axon targeting. Knocking down *Ptp10D* using a VA1d-ORN-specific-GAL4 driver^32^ and multiple RNAi lines caused similar mistargeting of VA1d-ORN axons to the VA1v glomerulus and mismatching with VA1v-PN dendrites (Fig. 2d, k; Extended Data Fig. 3a, b; Extended Data Table 2). Additional experiments argued against Ptp10D mediating homophilic attraction (Extended Data Fig. 3f, g). Furthermore, using a sparse VA1d-ORN GAL4 driver (Extended Data Fig. 4a–d) to knock down *Ptp10D* in single VA1d-ORN also caused axon branches to mistarget to the VA1v glomerulus (Fig. 2e, k), indicating that Ptp10D acts cell-autonomously in VA1d-ORNs to prevent their mismatching with VA1v-PNs.

Based on the inverse expression pattern of Ptp10D and Toll2 (Fig. 1h), we tested whether Toll2 plays a similar role in the precise targeting of VA1d-ORNs. Toll2 is a single-pass transmembrane protein belonging to the Toll-like receptor family, with leucine-rich repeats (LRRs) extracellularly and a Toll/interleukin-1 receptor (TIR) domain intracellularly (Fig. 2a). Toll2 plays an evolutionarily conserved roles in innate immunity and regulate tissue morphogenesis^44–46^, but its role in neural development is unclear. We found that both pan-PN RNAi-mediated knockdown of *Toll2* and loss of *Toll2* caused VA1d-ORN axons mistargeting to the VA1v glomerulus (Fig. 2f, k and Extended Data Fig. 1c).

Given the high expression of Toll2 in VA1v-PNs (Fig. 1h), we hypothesized that Toll2 in VA1v-PN dendrites sends a trans-cellular repulsive signal to VA1d-ORN axons to prevent misconnection between them. To manipulate Toll2 specifically in VA1v-PNs, we identified a VA1v-PN driver that labels VA1v-PNs across developmental stages (Extended Data Fig. 4e–i). Indeed, *Toll2* knockdown in VA1v-PNs caused VA1d-ORN axons mistargeting to the VA1v glomerulus (Fig. 2g, k), phenocopying *Ptp10D* knockdown in VA1d-ORNs. Thus, Toll2 acts in VA1v-PNs whereas Ptp10D acts in VA1d-ORNs to prevent misconnections between VA1d-ORN axons and VA1v-PN dendrites (Fig. 2m).

Since Ptp10D and Toll2 were also highly expressed in VA1d-PNs and VA1v-ORNs, respectively (Fig. 1h), we examined whether they are similarly required for preventing mismatching between VA1v-ORNs and VA1d-PNs. Indeed, cell-type-specific knockdown of *Ptp10D* in VA1d-PNs and Toll2 in VA1v-ORNs caused similar phenotypes: VA1d-PN dendrites mistargeted to the VA1v glomerulus (Fig. 2h–j, l; Extended Data Fig. 3c). Conversely, no mistargeting phenotype was observed in VA1d-or VA1v-ORN axons when *Toll2* was knocked down in ORNs (Extended Data Fig. 3d, e), suggesting that Toll2 does not function cell-autonomously in VA1d-or VA1v-ORNs and does not mediate interactions between VA1d-ORNs and VA1v-ORNs. No mistargeting phenotype was observed when knocking down *Toll2* or *Ptp10D* where they had low expression (Extended Data Table 2). Altogether, these data suggest that both Ptp10D and Toll2 act in both PNs and ORNs to prevent mismatching between non-partners, with Toll2 sending and Ptp10D receiving a repulsive signal to non-partner neurons (Fig. 2m).

### Trans-cellular interactions of Toll2 and Ptp10D

We next tested whether Toll2 and Ptp10D work together to prevent mismatching between non-partner PNs and ORNs via trans-cellular interactions. To do so, we first overexpressed Toll2 specifically in VA1d-ORNs, where the endogenous Toll2 level is low. This caused some of their partner VA1d-PN dendrites (with high Ptp10D) to mismatch with DA4l-ORN axons (with low Toll2) (Fig. 3a, b, e; Extended Data Fig. 5b); the same manipulation did not cause mistargeting phenotype in VA1d-ORN axons nor VA1v-PN dendrites (Extended Data Fig. 5a and Extended Data Table 2). This result supports the repulsion hypothesis: misexpressed Toll2 in VA1d-ORN axons sent a trans-cellular signal to repel the partner VA1d-PN dendrites away from them. Similarly, overexpressing Ptp10D specifically in VA1v-ORNs (whose synaptic partner VA1v-PNs expressed high Toll2) caused their axons to mismatch with VA1d-ORNs (with low Toll2; Extended Data Fig. 5c).

**Fig. 3.**
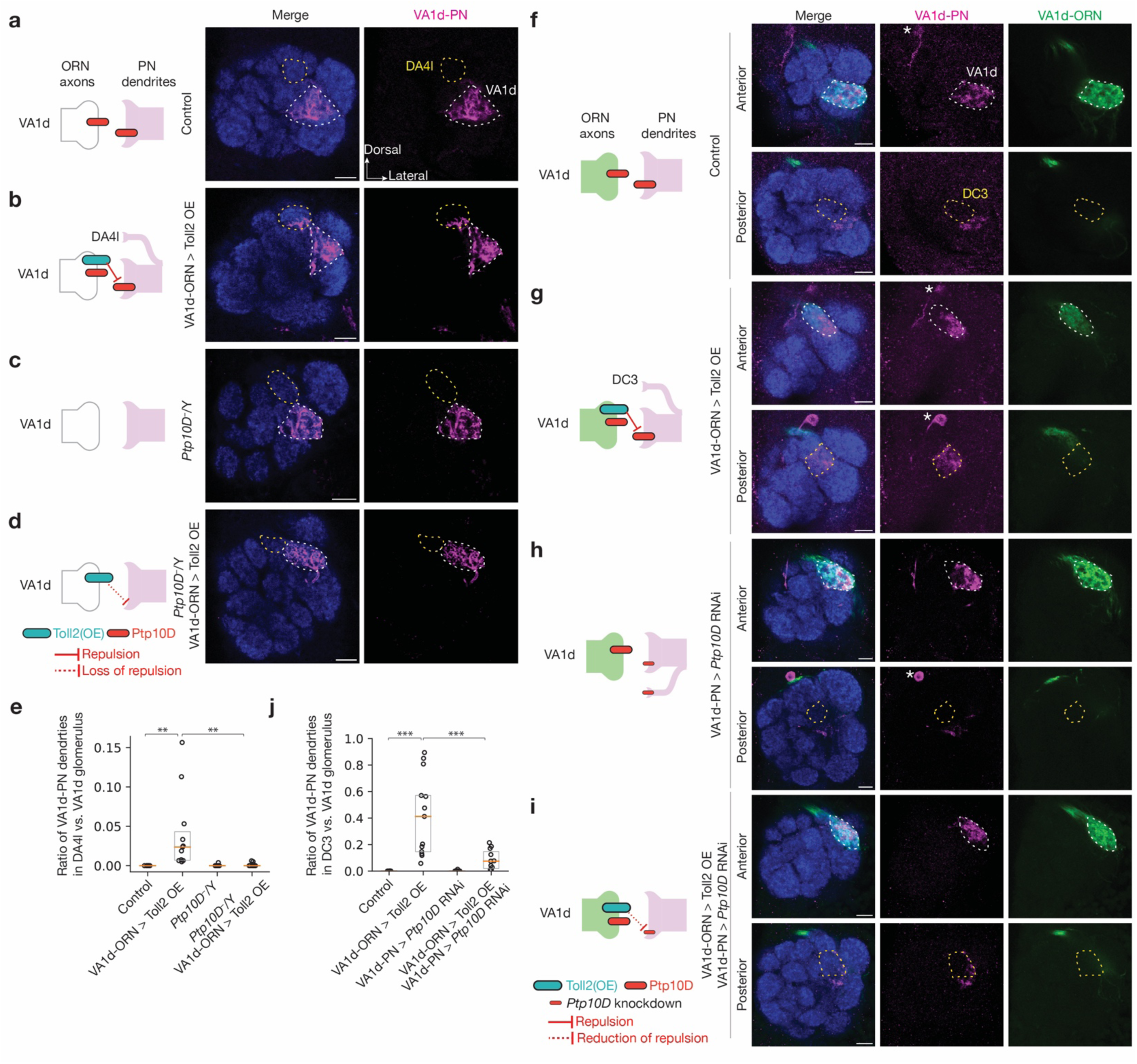
Ptp10D and Toll2 promote trans-cellular repulsive interactions. **a–d,** Left column shows experimental schematic. Middle and right columns are confocal images showing neuropil staining by N-cadherin antibody (blue) and VA1d-PN dendrites (magenta). The VA1d (white dashed lines) and DA4l (yellow dashed lines) glomeruli are outlined based on N-cadherin staining. Control VA1d-PN dendrites only innervate the VA1d glomerulus (**a**). Some VA1d-PN dendrites mistarget to the DA4l glomerulus following Toll2 overexpression in VA1d-ORNs (**b**). No VA1d-PN dendrites mistarget to the DA4l glomerulus in *Ptp10D* hemizygous mutant (**c**). Almost no VA1d-PN dendrites mistarget to the DA4l glomerulus when Toll2 overexpression in VA1d-ORNs was performed in *Ptp10D* hemizygous mutant (**d**). **e,** Quantification of mistargeting ratio of VA1d-PN dendrites in the DA4l versus VA1d glomerulus for **a–d**. **f–i,** Left column shows experimental schematic. Right three columns are confocal images showing neuropil staining by N-cadherin antibody (blue), VA1d-PN dendrites (magenta), and VA1d-ORN axons (green). The top and bottom rows are optical sections in the anterior and posterior parts of a representative antennal lobe, respectively. VA1d and DC3 glomeruli are outlined in white and yellow on the top and bottom row, respectively, based on N-cadherin staining. In control, VA1d-PN dendrites only innervate the VA1d glomerulus and fully overlap with VA1d-ORN axons (**f**). VA1d-PN dendrites overlap less with VA1d-ORN axons within the VA1d glomerulus and mistarget to the DC3 glomerulus following Toll2 overexpression in VA1d-ORNs (**g**). No VA1d-PN dendrites mistarget to the DC3 glomerulus when *Ptp10D* RNAi was expressed in VA1d-PNs (**h**). VA1d-PN dendrites overlap more with VA1d-ORN axons and mistarget less to DC3 glomerulus when Toll2 overexpression in VA1d-ORNs is combined with *Ptp10D* knockdown in VA1d-PNs (**i**). The different mistargeting regions in **b, e** and **g, j** likely result from different Toll2 overexpression levels using different binary systems (Methods). Both DA4l-ORNs and DC3-ORNs express low levels of Toll2, consistent with our repulsion model. * designates PN cell bodies. **j,** Quantification of the mistargeting ratio of VA1d-PN dendrites in the DC3 versus VA1d glomerulus for **f–i**. For all genotypes, n ≥ 10. Boxes in **e** and **j** indicate geometric mean and 25% to 75% range. Kruskal–Wallis test with Bonferroni’s multiple comparison.

Next, we tested whether the Toll2 repulsive signal was received by Ptp10D, which was highly expressed in VA1d-PN dendrites. We combined Toll2 overexpression in VA1d-ORNs with loss of Ptp10D. The mistargeting level of VA1d-PN dendrites to the DA4l glomerulus was significantly reduced in *Ptp10D* hemizygous mutant animals (Fig. 3d, e). *Ptp10D* hemizygous itself did not cause VA1d-PN dendrite mistargeting to DA4l, even though some VA1d-PN dendrites mistargeted to VA1v (Fig. 3c, e). This suppression indicates that Ptp10D is necessary to mediate the Toll2 overexpression phenotype, and thus Toll2 and Ptp10D function together to mediate repulsion.

As Ptp10D had high expression in both VA1d-PNs and VA1d-ORNs, the experiments above did not distinguish whether the suppression by Ptp10D knockout was a result of *cis-* or *trans*-interaction between Toll2 and Ptp10D, or a result of loss of Ptp10D in other glomeruli besides VA1d. To distinguish between these possibilities, we overexpressed Toll2 in VA1d-ORNs and knocked down *Ptp10D* in VA1d-PNs simultaneously using two orthogonal binary expression systems. In wild-type flies, dually labeled VA1d-ORN axons and VA1d-PN dendrites largely intermingled with each other (Fig. 3f, j). Overexpressing Toll2 in VA1d-ORNs caused VA1d-PN dendrites to segregate from VA1d-ORN axons within the VA1d glomerulus and mistarget to the nearby DC3 glomerulus (Fig. 3g, j). Simultaneous knockdown of *Ptp10D* in VA1d-PNs and overexpression of Toll2 in VA1d-ORNs suppressed the VA1d-PN dendrite phenotypes caused by Toll2 overexpression alone (Fig. 3i, j), whereas *Ptp10D* knockdown in VA1d-PNs alone did not cause a similar phenotype (Fig. 3h, j). Taken together with the inverse expression of Toll2 and Ptp10D and their similar loss-of-function phenotypes, these trans-cellular interaction data support a model in which Toll2 sends and Ptp10D receives a repulsive signal to prevent matching between non-partner ORNs and PNs (Fig. 2m).

### Fili and Kek1 for non-partner PN-ORN repulsion

To study the function of the other CSP pairs, we performed similar loss-of-function and suppression experiments as with Toll2–Ptp10D. A previous study showed that when *Fili* is knocked out or knocked down in VA1d/DC3-PNs, VA1v-ORN axons mistarget to the VA1d glomerulus whereas VA1d-ORN axons and VA1d/DC3-PN dendrites are unaffected. This result suggests that Fili is required in VA1d/DC3-PNs to prevent mistargeting of VA1v-ORN axons to the VA1d glomerulus^27^. However, Fili’s CSP partner remained elusive. Kek1 was a top candidate based on its high expression in VA1v-ORNs (Fig. 1i; Extended Data Fig. 2) and mistargeting of VA1v-ORN axons to the VA1d glomerulus caused by pan-ORN knockdown of *kek1* (Extended Data Fig. 1e). Kek1 and Fili both contain leucine-rich repeats (LRRs) in their extracellular domain (Fig. 4a). Kek1 inhibits epidermal growth factor receptor (EGFR) activity through the LRRs during eye development^47^ and is expressed in developing CNS with poorly defined function^48^. We found that both homozygous mutant of *kek1* and *kek1* knockdown in VA1v-ORNs caused VA1v-ORN axons to mistarget to the VA1d glomerulus (Fig. 4b–e), phenocopying the loss of Fili in VA1d/DC3 PNs^27^.

**Fig. 4.**
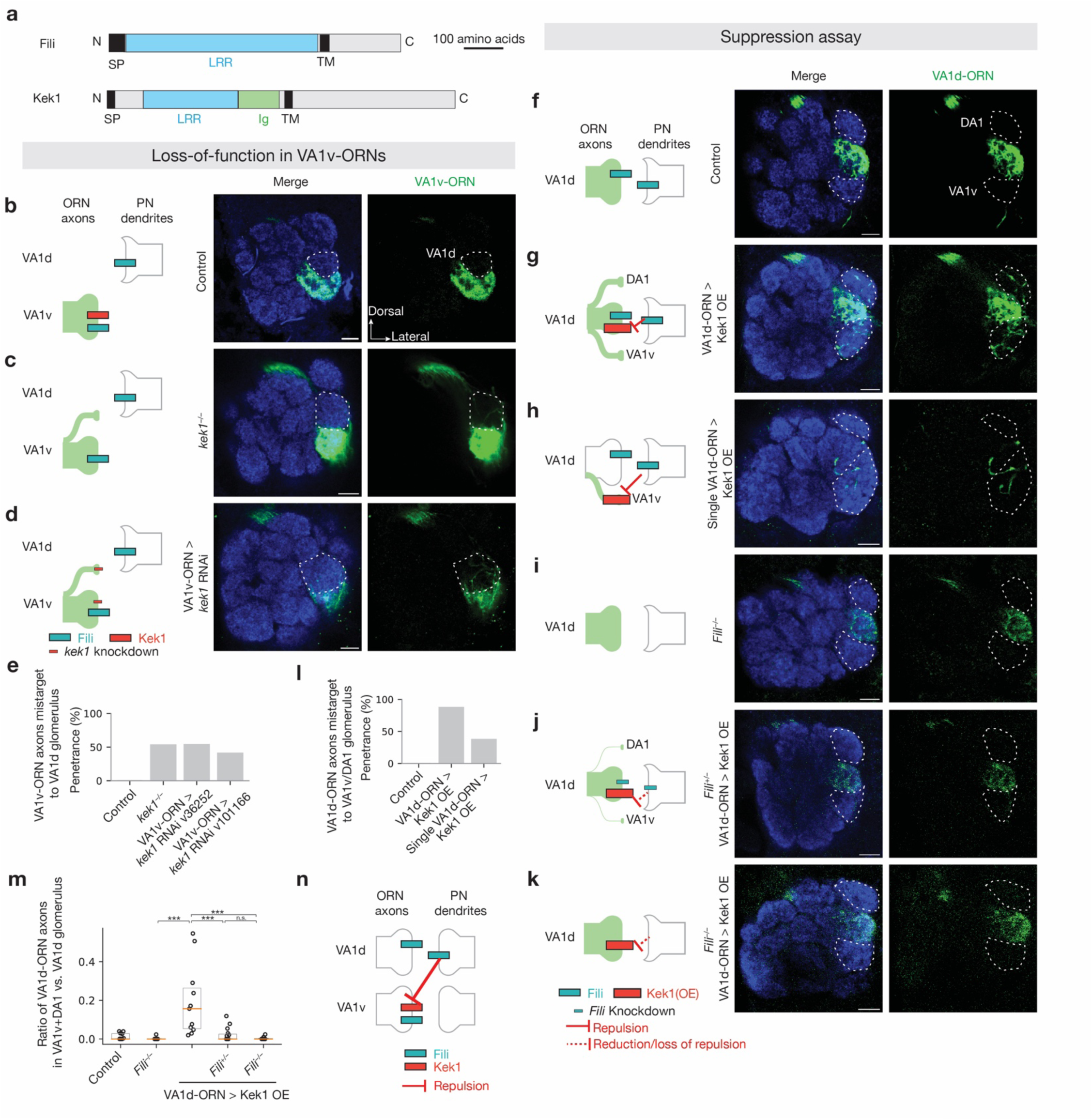
Repulsive interactions between Fili and Kek1. **a,** Domain composition of *Drosophila* CSPs Fili and Kek1. TM, transmembrane domain; N, N-terminus; C, C-terminus; SP, signal peptide; LRR, leucine-rich repeat; Ig, immunoglobulin domain. **b–d,** Left column shows experimental schematic. Middle and right columns are confocal images showing adult antennal lobe neuropil staining by N-cadherin antibody (blue) and VA1v-ORN axons (green). The VA1d glomerulus is outlined based on N-cadherin staining. Control VA1v-ORN axons only innervate the VA1v glomerulus ventral to the VA1d glomerulus (**b**). Some VA1v-ORN axons mistarget to the VA1d glomerulus in *kek1* mutant (**c**) or *kek1* RNAi expressed in VA1v-ORNs (**d**). **e,** Penetrance of the mistargeting phenotypes in **b–d**. For all genotypes, n ≥ 10. **f–k,** Left column shows experimental schematic. Middle and right columns are confocal images showing neuropil staining by N-cadherin antibody (blue) and VA1d-ORN axons (green). DA1 and VA1v glomeruli are outlined based on N-cadherin staining. Control VA1d-ORN axons only innervate the VA1d glomerulus (**f**). Some VA1d-ORN axons mistarget to the DA1 and VA1v glomeruli following Kek1 overexpression in all (**g**) or single (**h**) VA1d-ORNs. No VA1d-ORN axons mistarget to the DA1 or VA1v glomeruli in *Fili* mutant (**i**). Almost no VA1d-ORN axons mistarget to the DA1 and VA1v glomeruli when Kek1 overexpression in VA1d-ORNs was performed in *Fili* heterozygous (**j**) or homozygous (**k**) mutant. **l,** Penetrance of the mistargeting phenotypes in **f–h**. For all genotypes, n ≥ 10. **m,** Quantification of the mistargeting ratio of VA1d-ORN axons in the DA1+VA1v versus VA1d glomerulus. For all genotypes, n ≥ 10. Boxes indicate geometric mean and 25% to 75% range. Kruskal–Wallis test with Bonferroni’s multiple comparison. **n,** Schematic summary for the function of Kek1 and Fili in the VA1d and VA1v glomeruli.

To further investigate whether Fili and Kek1 work together to prevent misconnections between non-partner PNs and ORNs, we overexpressed Kek1 specifically in VA1d-ORNs, whose synaptic partner VA1d-PNs expressed high Fili^27^. This caused VA1d-ORN axons to mistarget to the neighboring VA1v and DA1 glomeruli (Fig. 4f, g, m), whose PN dendrites mostly do not express Fili^27^. To test whether the Kek1 overexpression phenotype was caused by its interaction with Fili, we overexpressed Kek1 in *Fili* mutant animals (Fig. 4i, m). The overexpression phenotype was much less severe in *Fili* heterozygous mutant (Fig. 4j, m) and was nearly fully suppressed in *Fili* homozygous mutant (Fig. 4k, m). In addition, overexpressing Kek1 in single VA1d-ORN using the sparse driver produced a similar mistargeting phenotype (Fig. 4h, l), indicating that Kek1 acts cell-autonomously. Although VA1d-ORNs and VA1v-ORNs also expressed Fili (Fig. 1i), a previous study showed that Fili knockout in ORNs does not cause any mistargeting phenotype of VA1d-and VA1v-ORN axons^27^. Together, these data suggest that Fili and Kek1 work together to prevent the misconnections between non-partner PNs and ORNs, with Fili sending and Kek1 receiving the trans-cellular repulsive signal (Fig. 4n).

### Hbs/Sns and Kirre for non-partner PN-ORN repulsion

Hbs/Sns and Kirre are evolutionarily conserved immunoglobulin family of ligand-receptor pairs with conserved binding sites^49^ (Fig. 5a). *Drosophila* Hbs/Sns and Kirre regulate myoblast fusion^50^, nephrocytes functions^51^, and neural circuit wiring^52^. Whereas previous studies have mostly suggested that this ligand-receptor pair (and sometimes homophilic interactions) mediates attraction^53,54^, the expression patterns in the fly olfactory system (Fig. 1j) and RNAi phenotypes (Extended Fig. 1f–h) raised the possibility that Hbs/Sns and Kirre might also mediate repulsion. We found that *kirre*, *hbs*, or *sns* mutants, as well as VA1v-ORN-specific knockdown of *kirre* and VA1d-PN-specific knockdown of *hbs* or *sns*, all caused a similar phenotype: mistargeting of VA1v-ORNs to the VA1d glomerulus (Fig. 5b–i). Thus, Kirre, Hbs, and Sns are all required to prevent VA1v-ORNs from matching with VA1d-PNs, with Kirre acting in VA1v-ORNs and Hbs/Sns acting in VA1d-PNs to prevent VA1v-ORNs to mistarget to the VA1d glomerulus.

**Fig. 5.**
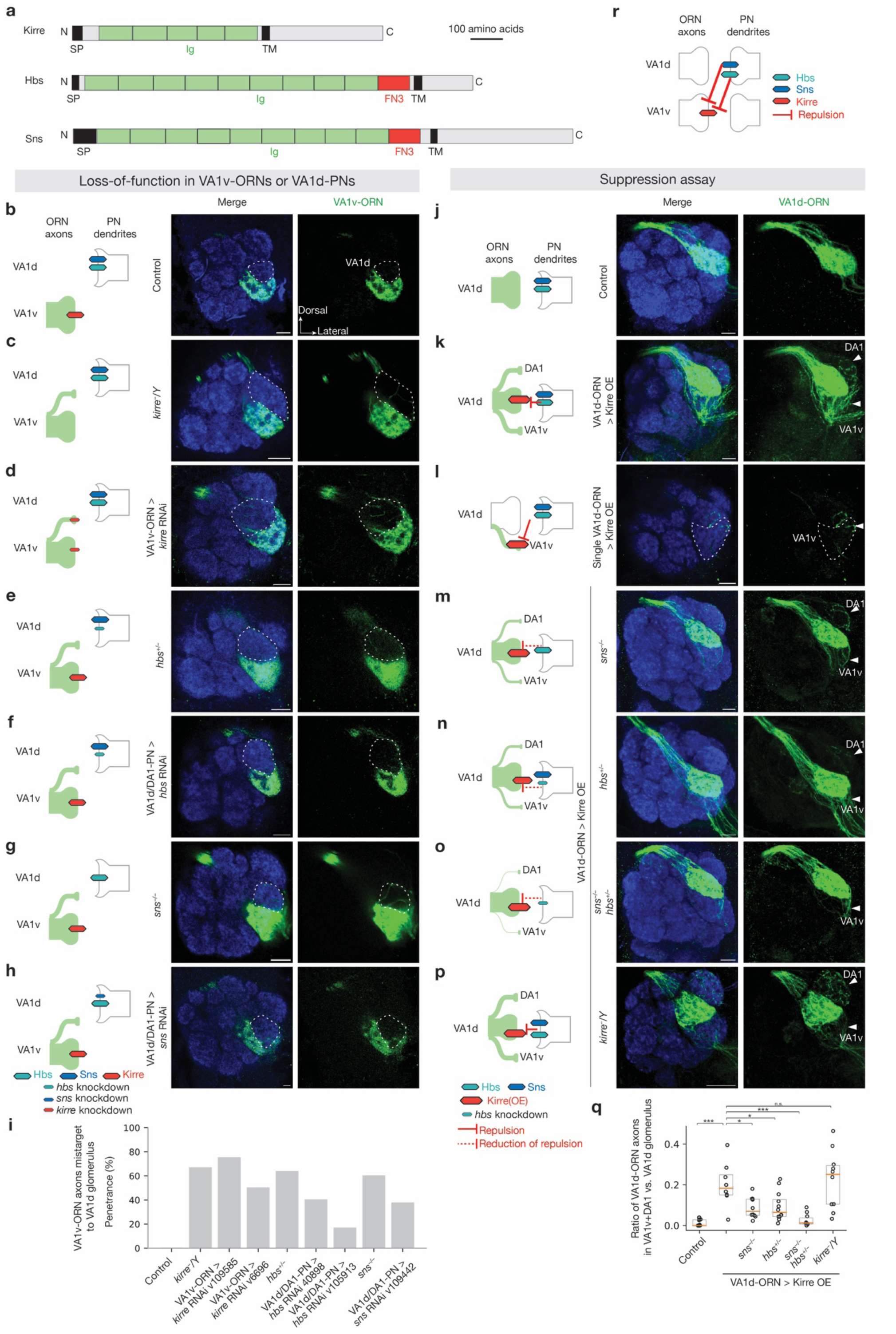
Repulsive interactions between Kirre and Hbs/Sns. **a,** Domain composition of *Drosophila* cell-surface proteins Kirre, Hbs, and Sns. TM, transmembrane domain; N, N-terminus; C, C-terminus; SP, signal peptide; Ig, immunoglobulin domain; FN3, fibronectin type III domain. **b–h,** Left column shows experimental schematic. Middle and right columns are confocal images of adult antennal lobes showing neuropil staining by N-cadherin antibody (blue) and VA1v-ORN axons (green). The VA1d glomerulus is outlined based on N-cadherin staining. Control VA1v-ORN axons only innervate the VA1v glomerulus (**b**). Some VA1v-ORN axons mistarget to the VA1d glomerulus in *kirre* hemizygous mutant (**c**), *kirre* RNAi expressed in VA1v-ORNs (**d**), *hbs* heterozygous mutant (**e**), *hbs* RNAi expressed in DA1-PNs and VA1d-PNs (**f**), *sns* homozygous mutant (**g**), or *sns* RNAi expressed in DA1-PNs and VA1d-PNs (**h**). **i,** Penetrance of the mistargeting phenotypes in **b–h**. For all genotypes, n ≥ 10. **j–p,** Left column shows experimental schematic. Middle and right columns are maximum projections (**j–k, m–p**) or a single section (**l**) of confocal images showing neuropil staining by N-cadherin antibody (blue) and VA1d-ORN axons (green). Arrowheads indicate the DA1 and VA1v glomeruli based on N-cadherin staining. Control VA1d-ORN axons only innervate the VA1d glomerulus ventral to the DA1 glomerulus and dorsal to the VA1v glomerulus (**j**). Some VA1d-ORN axons mistarget to the DA1 and VA1v glomeruli following Kirre overexpression in all (**k**) or single (**l**) VA1d-ORNs. Comparing to Kirre overexpression alone, fewer VA1d-ORN axons mistarget to the DA1 and VA1v glomeruli when Kirre overexpression in VA1d-ORNs was performed in *sns* homozygous mutant (**m**), *hbs* heterozygous mutant (**n**), or the combination of them (**o**). VA1d-ORN axons still mistarget to the DA1 and VA1v glomerulus when Kirre overexpression in VA1d-ORNs was performed in *kirre* hemizygous mutant (**p**). **q,** Quantification of the mistargeting ratio of VA1d-ORN axons in the DA1+VA1v versus VA1d glomerulus. For all genotypes, n ≥ 10. Boxes indicate geometric mean and 25% to 75% range. Kruskal–Wallis test with Bonferroni’s multiple comparison. **r,** Schematic summary for the function of Kirre, Hbs, and Sns in ORN-PN matching in the VA1d and VA1v glomeruli.

To examine whether Hbs/Sns and Kirre work together to instruct synaptic partner matching, we performed genetic interaction experiments. We first overexpressed Kirre in VA1d-ORNs, whose partner, VA1d-PNs, expressed high Hbs and Sns (Fig. 1j). Kirre overexpression in all VA1d-ORNs or single VA1d-ORN led to mistargeting of some VA1d-ORN axons to the VA1v and DA1 glomeruli (Fig. 5j–l, q), whose PNs expressed low Hbs and Sns, suggesting cell-autonomous function of Kirre as a repulsive receptor. Overexpressing Kirre in *hbs* mutant, *sns* mutant, or *hbs*/*sns* double mutant background reduced mistargeting phenotypes of VA1d-ORN axons (Fig. 5m–o, q). None of the mutants alone had any VA1d-ORN axon mistargeting phenotype in the absence of Kirre overexpression (Extended Data Table 2). Since Kirre can also mediate homophilic binding^49^, we tested whether Kirre homophilic attraction is responsible for the mistargeting phenotype we observed. Overexpressing Kirre specifically in VA1d-ORNs caused the same phenotype in *kirre* hemizygous mutant animals as in wild type (Fig. 5p, q), arguing against a contribution for trans-cellular Kirre homophilic interaction in synaptic partner matching. Together, these data support that heterophilic repulsion between Hbs/Sns as the ligands and Kirre as the receptor prevents VA1v-ORNs from mistargeting to the VA1d glomerulus (Fig. 5r).

As the genetic interactions of Fili–Kek1 and Hbs/Sns–Kirre both functions to prevent misconnection of VA1v-ORNs with VA1d-PNs, we tested whether there is crosstalk between these interactions. We found that knocking down *sns* and *hbs* did not suppress the mistargeting phenotype caused by Kek1 overexpression (Extended Data Fig. 6), suggesting that Fili–Kek1 and Hbs/Sns–Kirre likely act in distinct pathways and validating the specificity of these genetic suppression experiments. In an additional set of experiments, we co-expressed Bruchpilot-Short, a presynaptic active zone marker, and found that it was enriched in mistargeted ORN axons when we perturbed each of the three CSP pairs (Extended Data Fig. 7). These data suggest that mistargeted ORN axons may form ectopic synaptic connections with new PN partners.

As previous biochemical and structural studies have shown direct binding between Hbs/Sns and Kirre^49^, we also tested whether the other two CSP pairs we identified directly bind each other. However, we did not detect direct binding between Fili and Kek1, or between Toll2 and Ptp10D, in *in vitro* (Extended Data Fig. 8) or tissue-based (Extended Data Fig. 9) binding assays. Thus, the biochemical basis for the repulsive genetic interactions mediated by Fili–Kek1 and Toll2–Ptp10D remains an open question. Possibilities include requirements for unidentified co-factor(s), post-translational modifications, or specific physiological conditions like multimerization^55,56^ that were not recapitulated in our binding assays.

### Broad usage of the three CSP pairs

Our results (Figs. 2–5) and additional control experiments (Extended Data Table 2) suggest that the three repulsive CSP pairs could prevent mismatching between VA1d-ORNs with VA1v-PNs (Toll2–Ptp10D), VA1d-PNs with VA1v-ORNs (Toll2–Ptp10D), and VA1v-ORNs with VA1d-PNs (Fili–Kek1; Hbs/Sns–Kirre) (Fig. 6a). Thus, the three CSP pairs work in concert, with partial redundancy, to ensure robust repulsion between non-matching ORNs and PNs at the VA1d and VA1v glomeruli.

**Fig. 6.**
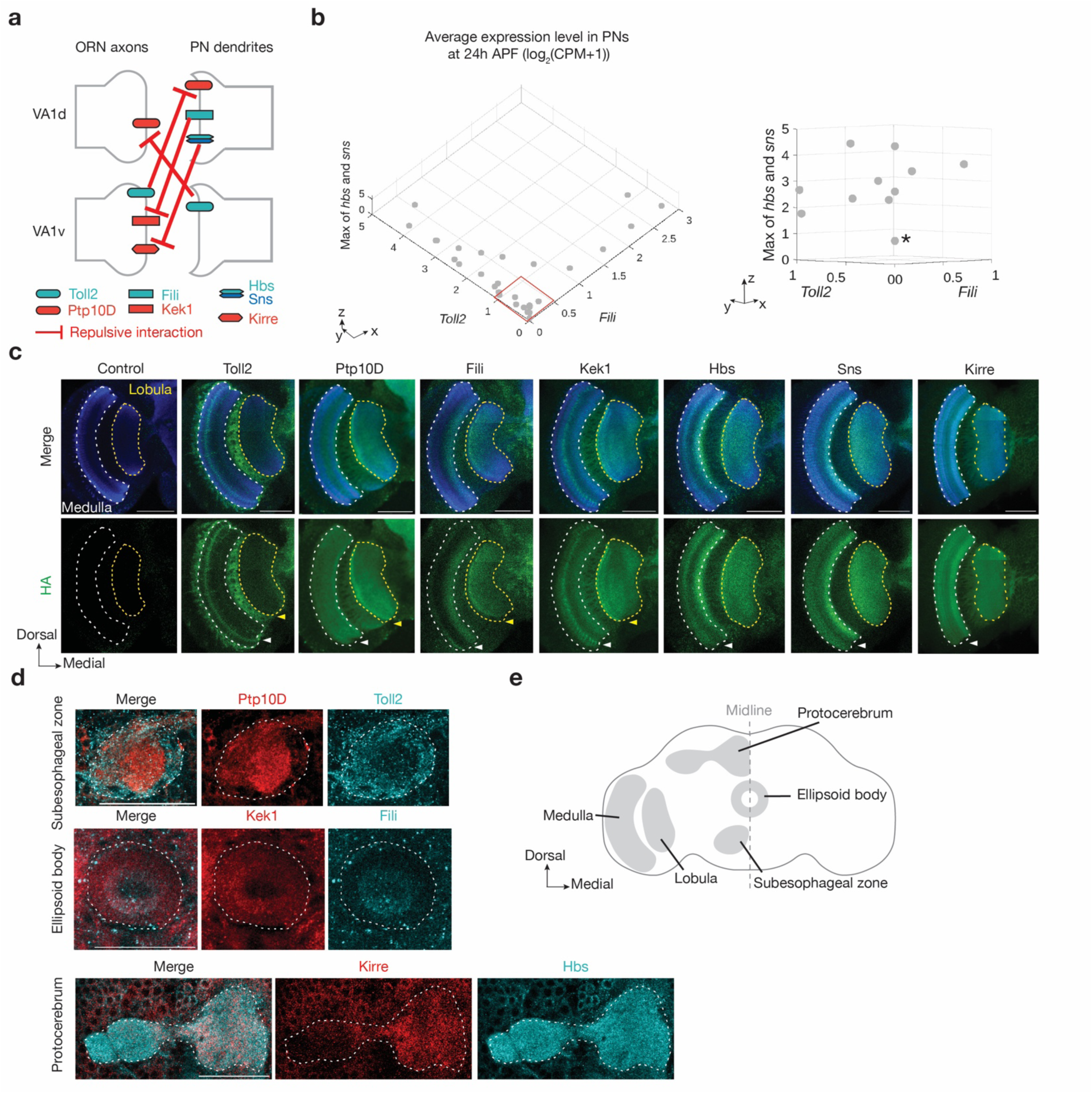
Three CSP pairs collectively cover the antennal lobe and are differentially expressed in other brain regions. **a,** Schematic summary of repulsive interactions of three CSP pairs in the VA1v and VA1d glomeruli. Hbs and Sns are combined as they perform similar functions. **b,** Single-cell RNA sequencing data showing the average expression level of non-cell-autonomous cues *Toll2*, *Fili,* and the maximum of *hbs* and *sns* in PNs at 24–30h APF. Right column is a zoom-in of the red box in the left column from a different angle, showing that PN types that express low levels of *Toll2* and *Fili* tend to express high levels of *hbs* or *sns* (* indicates a possible exception). Units for all axis: log_2_(CPM+1). CPM: counts per million reads. Data adapted from previous published studies^3,4^. **c,** Confocal images showing neuropil staining by N-cadherin antibody (blue) and HA staining (green) of tagged endogenous cell-surface proteins in the *Drosophila* optic lobe at 42–48 h APF. The medulla (white dotted lines) and lobula (yellow dotted lines) are outlined based on N-cadherin staining. Arrowheads indicate the layers with differential expression patterns. For example, Kek1 is highly expressed in a medulla layer that is low for Fili expression. Scale bars = 50 µm. **d,** Confocal images showing HA staining (for Ptp10D, Kek1, and Hbs) and V5 staining (for Toll2, Fili, and Kirre) of tagged endogenous CSP pairs in the same brain (each row) at 42–48 h APF. The subesophageal zone, ellipsoid body, and protocerebrum regions are outlined based on neuropil staining. Scale bars = 50 µm. **e,** Pupal *Drosophila* brain schematic with brain regions in **c** and **d** highlighted in grey in the left hemisphere.

The antennal lobe has 50 ORN-PN synaptic partner pairs that need to be specified, potentially requiring many CSP pairs to mediate repulsive interactions. One way to alleviate this is to repeatedly use the same repulsive CSP pairs across the antennal lobe in a combinatorial fashion. Indeed, analysis of previously published single-cell transcriptomes during development^3^ revealed that mRNAs encoding all three CSPs implicated in sending the repulsive cues—*Toll2*, *Fili*, and the maximum of *hbs* and *sns*—are expressed in multiple PN types. Interestingly, most PN types expressed one or two but not all the three repulsive cues at high levels (Fig. 6b). For each PN type, this property reduces the errors of mismatching with non-partner ORN types (as all PN types express at least one repulsive cue at high levels) while making room for the match with its partner ORN type (as few PN type expresses all three repulsive cues at high levels).

Based on the expression pattern, we further investigated whether the repulsive interactions of these three CSP pairs play similar roles in additional ORN and PN types using loss-or gain-of-function experiments. A previous study showed that Fili acts in ORNs to prevent mistargeting of VM5-PN dendrites^27^. We observed a similar mistargeting phenotype of VM5-PN dendrites when we knocked down *kek1* using a pan-PN driver (Extended Data Fig. 10a, b), suggesting that Fili in ORNs and Kek1 in PNs are both required for proper targeting of VM5-PN dendrites. As Hbs is highly expressed in DA1-PNs and DA4l-PNs, we overexpressed Kirre in DA1-ORNs or DA4l/VA1d-ORNs using available drivers with early developmental onset. We observed mistargeting of each ORN group(s) to neighboring glomeruli, whose PNs expressed low Hbs and Sns (Extended Data Fig. 10c-e), suggesting that Hbs and Kirre mediate repulsive interactions in these PN-ORN pairs. In the companion manuscript, we show that all the three repulsive CSP pairs play a key role in preventing the mismatch of both VA1d-ORNs and DA1-ORNs with PNs of nearby glomeruli, as manipulating the expression of these CSP pairs is essential in the rewiring of VA1d-and DA1-ORNs to non-cognate PNs^37^. Altogether, these data support a model that these repulsive interactions are used broadly in the antennal lobe to ensure synaptic partner matching specificity.

Finally, to explore the possibility that these three CSP pairs work elsewhere in the fly brain to regulate connection specificity, we used HA staining to examine the expression patterns of the 7 CSPs (Fig. 1b) during the period when many circuits are establishing wiring specificity (42–48h APF). We found that all CSPs had broad but differential expression across the *Drosophila* brain (Supplementary Videos 1–7). For example, in the medulla and lobula of the optic lobe, all CSPs showed differential expressions in specific neuropil layers (Fig. 6c, e). To better examine the endogenous expression pattern of CSP pairs in brain regions without layer structures, we also produced transgenic flies with endogenous CSPs tagged with V5 and co-labeled each pair of CSPs in the same brain. Fig. 6d showcased the differential expression of these CSP pairs in the subesophageal zone, ellipsoid body, and protocerebrum. These data support the notion that combinatorial repulsive interactions serve as a generalizable mechanism in instructing wiring specificity of neural circuits.

## Discussion

Previous high-throughput extracellular interactome screenings *in vitro* have identified novel molecular pairs with direct interactions, including Dpr/DIP and Beat/Side families of immunoglobulin-containing CSPs whose *in vivo* functions in neuronal wiring have subsequently been validated^15–17,20–22,57,58^. Here, we took an alternative approach of using transcriptome-informed *in vivo* genetic screens, which enabled us to identify known binding partners (Hbs/Sns–Kirre) as well as proteins that may not interact directly (Fili–Kek1 and Toll2–Ptp10D). Thus, transcriptome-informed *in vivo* screening complements *in vitro* biochemical approach to identify new CSP pairs that mediate trans-cellular interactions in neuronal wiring.

The inverse expression patterns of the three CSP pairs we identified, their cell-type-specific loss-of-function phenotypes, and suppression assays strongly suggest that they mediate repulsion between non-partner PNs and ORNs. Some of the CSPs only have modest differential expression, suggesting that synaptic partner choice is likely regulated by the relative levels of repulsive interactions. We note that orthologs of Hbs/Sns and Kirre in other species control synaptic site choice in *C. elegans*^53^ and axon sorting in the mouse olfactory bulb^54^ via heterophilic and homophilic attraction, respectively. However, our results suggest they instruct synaptic partner matching in the fly olfactory system via heterophilic repulsion. These different mechanisms could potentially be mediated by engaging distinct intracellular signaling pathways in specific cellular context.

Repulsion could be combined with attraction to enhance the selection process of synaptic partners. For example, for neuron A to match its synaptic partner A’ but not non-partners, one strategy is to express attractive CSP pairs in A and A’. However, as the CSP number on each synaptic partner increases, an attraction-only strategy can cause ambiguity (for example, to distinguish the matching of two vs. three attractive pairs) and the addition of repulsion can reduce errors. Repulsion can also increase the searching efficiency by ruling out non-partners during the simultaneous searching process, as in the case of ORN-PN matching^31,32^. On the other hand, a repulsion-only strategy may have difficulty exploring a larger space due to excessive branch retraction. In the companion manuscript, we showed that only by simultaneously manipulating both attractive and repulsive CSPs in ORNs could we substantially switch their partner PNs^37^. Thus, attractive and repulsive interactions work in concert to ensure precise synaptic partner matching.

## Methods

### Generation of cDNA constructs and transgenic flies

Complementary DNA (cDNA) encoding for proteins used in this study were obtained from different resources. *Toll2* cDNA was amplified from the cDNA library of *w1118* pupal brain extracts using Q5 hot-start high-fidelity DNA polymerase (New England Biolabs) as previously described^32^; *Ptp10D* cDNA was amplified from clone RE52018 (DGRC Stock 9073; https://dgrc.bio.indiana.edu//stock/9073; RRID:DGRC_9073); *Fili* cDNA was amplified from *pUAST-attB-SP-V5-Fili-FLAG* plasmid^27^; *kek1* cDNA was amplified from clone GH23277 (DGRC Stock 1263019; https://dgrc.bio.indiana.edu//stock/1263019; RRID:DGRC_1263019); *kirre* cDNA was amplified from genomic DNA extraction of *UAS-kirre.C-HA* fly^59^ (RRID:BDSC_92196) using DNeasy blood and tissue kit (QIAGEN). Sequence-verified coding regions were assembled into *pUAST-attB-mtdT-3xHA*^60^, *pUAST-attB-SP-V5-Fili-FLAG*^32^ or *pJFRC19-13XLexAop2-IVS-myr::GFP* (Addgene Plasmid #26224) backbones using the NEBuilder HiFi DNA assembly master mix (New England Biolabs) to generate *pUAST-attB-Toll2-FLAG, pUAST-attB-Ptp10D-3xHA, pUAST-attB-kek1-3xHA, pUASTattB-kirre-3xHA,* and *pJFRC19-13XLexAop2-IVS-Toll2* plasmids. Transgenic flies for overexpression experiments were generated by BestGene with microinjection of plasmids into the *VK5* site.

### Generation of conditional tags

Endogenous conditional tag flies were generated using CRISPR knock-in with modifications from a previous strategy^38^. To increase the efficiency of knock-in, we incorporated the short repair templates flanked by gRNA target sites^61^ and the gRNA into a single plasmid *TOPO-HR1-FRT-3xHA-Stop-FRT-3xMyc-Stop-loxP-mCherry-loxP-HR2-pU6-gRNA* or *TOPO-HR1-FRT-V5-Stop-FRT-Flag-Stop-loxP-mCherry-loxP-HR2-pU6-gRNA*. In the plasmid, *HR1* and *HR2* are the 150-bp genomic sequences of upstream and downstream of the target genes stop codon, respectively; *gRNA* sequences were designed by the flyCRISPR Target Finder tool that targeting stop codons of the genes, and were cloned into the backbone of *pU6-BbsI-chiRNA* vector^62,63^ to make the *pU6-gRNA*. The plasmids were synthesized by Synbio Technologies and were microinjected in house into *nos-Cas9* flies^64^. All *mCherry*+ progenies were individually balanced and the *loxP*-flanked *mCherry* cassettes were then removed by crossing each line to balancer expressing Cre (Bloomington *Drosophila* Stock Center, RRID:BDSC 1092). To detect cell-type-specific expression level, we used *ey-Flp*^65^ for ORNs and *VT033006-GAL4;UAS-FLP*^66^ for PNs.

### Generation of the sparse driver and single-neuron genetic manipulations

The *FRT100-Stop-FRT100* element was cloned into the *78H05-p65AD* plasmid (in backbone *pBPp65ADZpUw*) to generate plasmid *78H05-FRT100-Stop-FRT100-p65AD* as previously reported^67^. The plasmid was integrated into the *VK27* site. To perform the sparse genetic manipulations, flies including VA1d-ORN sparse driver (*31F09-GAL4DBD*, *78H05-FRT100-Stop-FRT100-p65AD*), *hsFLP* (heat shock protein promoter-driven FLP), reporter (*UAS-mCD8-GFP*), and knockdown/overexpression transgenes were raised at 29°C. To induced sparse manipulation, the flies were collected at 0–6 hours after puparium formation and heat shocked for 1–2 hours in 37-degree water bath^67^.

### Immunostaining

Fly brains dissection, fixation, and immunostaining were done according to the published protocol^68^. For primary antibodies, we used rat anti-NCadherin (1:40; DN-Ex#8, Developmental Studies Hybridoma Bank), chicken anti-GFP (1:1000; GFP-1020, Aves Labs), rabbit anti-DsRed (1:500; 632496, Clontech), mouse anti-rat CD2 (1:200; OX-34, Bio-Rad), rabbit anti-HA (1:100, 3724S, Cell Signaling), mouse anti-HA (1:100, 2367S, Cell Signaling), rabbit anti-Myc (1:250, 2278S, Cell Signaling), and mouse anti-V5 (1:250, R960-25, Thermo Fisher Scientific). Donkey secondary antibodies conjugated to Alexa Fluor 405/488/568/647 or Cy3 (Jackson ImmunoResearch or Thermo Fisher) were used at 1:250. For the staining of conditional tag for Hbs and Fili in PNs, the routine protocol described above failed to detect Myc signal from the background, likely due to low expression of endogenous proteins *in vivo*. Alexa 488 Tyramide SuperBoost kit (Thermo Fisher) was used to amplify the immunostaining signal by following the manufacture’s protocol.

### Imaging, quantification, and statistical analysis

Images were obtained using laser scanning confocal microscopy (Zeiss LSM 780 or LSM 900). Fiji was used to adjust brightness and contrast for representative images. Penetrance of phenotypes represents the percentage of antennal lobes showing a given phenotype among the total antennal lobes (two per animal) examined. To quantify the endogenous expression levels of the proteins, we manually outlined VA1d and VA1v glomeruli in Fiji based only on the NCad signal (i.e., blind to Myc signals), and use this as filter to calculate mean fluorescent density 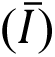 in the VA1d and VA1v glomeruli (total fluorescence intensity divided by the volume). The preference index is calculated in Python by 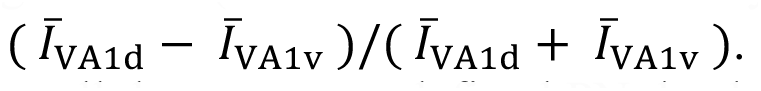 To quantify the mistargeting ratio of VA1d-PNs or VA1d-ORNs in the trans-cellular assay, we defined PN dendritic or ORN axonal targeting area by smoothening (‘gaussian blur’ with radius = 1 pixels) and thresholding (based on the algorithm ‘Otsu’) the images in Fiji. We manually outlined VA1d, VA1v, DA1, DA4l, and/or DC3 glomeruli in Fiji based only on the NCad signal (i.e., blind to PNs or ORNs signals), and used this as a filter to calculate PN dendritic or ORN axonal targeting volume (*V*) in each glomerulus. The mistargeting ratio is calculated in Python by 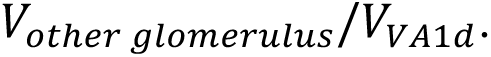

### Binding assays

For binding assays involving purified proteins with surface plasmon resonance, we expressed and purified Fili, Kek1, Ptp10D, and Toll2 extracellular domains using the baculoviral expression system in *Trichoplusia ni* High Five cells. Proteins were tagged with C-terminal hexahistidine tags for purification, and Avi-tags for biotinylation using BirA biotin ligase. Proteins were purified to homogeneity with Ni-NTA metal affinity and size-exclusion chromatography in 10 mM HEPES pH 7.2, 150 mM NaCl. Biotinylated Fili and Toll2 extracellular domains were captured on a Streptavidin sensor chip in a Biacore T200 system (Cytiva) running a buffer containing 10 mM HEPES pH 7.2, 150 mM NaCl and 0.05% Tween-20. We observed no binding responses for Kek1 and Ptp10D extracellular domains flowing on Fili and Toll2 channels, respectively.

For the avidity-based extracellular binding assay (ECIA), we followed the protocols published^69^. Briefly, we expressed and secreted each ectodomain as bait and/or prey, where they were tagged with Fc, V5, and His tags (for bait) and Alkaline phosphatase, the COMP pentameric coiled coil region, FLAG, and His tags (for prey). All proteins were expressed in *Drosophila* S2 cells in Schneider’s medium supplemented with the insect medium supplement (Sigma). Media from bait-expressing cells were incubated with Protein A-coated 96-well plates for capturing of bait onto plates overnight at room temperature, while media from prey-expressing cells were incubated with bait-captured plates for 3 hours. Washes were performed with 1x phosphate-buffered saline, supplemented with 1 mM CaCl_2_, 1 mM MgCl_2_ and 0.1% bovine serum albumin. Binding of prey to bait was detected using the chromogenic alkaline phosphatase substrate BluePhos (KPL) by measuring absorbance at 650 nm after a two-hour incubation. We included the known interactions of Rst dimerization^49^ and EGFR-Kek1 interaction as positive controls^70^.

The tissue-based binding assays were performed as previously described^58,71,72^. In brief, brains or wing discs were dissected in the Schneider’s medium, then incubated with the conditioned medium of High Five cells expressing epitope-tagged extracellular domains of a specific protein for 18 hours (for pupal brains) or 1 hour (for wing discs) at 4°C on a rotating platform. Medium of High Five cells without expressing any transgenes was used as a negative control. After the incubation, brains or wing discs were washed with the Schneider’s medium and fixed with 4% paraformaldehyde in 1x PBS for 30 minutes, followed by the immunostaining protocol above using antibodies against the epitope tags.

## Data and materials availability

All data are included in the manuscript, the supplementary materials, or are available upon request.

## Code availability

Code are available on github (https://github.com/ZhuoranLi97/repulsive_interactions).

## Acknowledgement

We thank Claude Desplan and Yu-Chieh David Chen for sharing split-GAL4 drivers; Mary Baylies for sharing *hbs/sns* double mutant flies; Mar Ruiz for sharing *kirre* mutant flies; Bloomington *Drosophila* Stock Center (BDRC), the Vienna *Drosophila* Resource Center (VDRC), and Fly Stocks of National Institute of Genetics (NIG) for providing fly lines; *Drosophila* Genomics Resource Center (DGRC) supported by NIH grant 2P40OD010949 and Addgene for providing plasmids. We also thank all the Luo laboratory members, especially Tom Hindmarsh Sten, Daniel Pederick, Colleen McLaughlin, Kin Lam Wong, Yunming Wu, Qijing Xie, Ji Hui, Jordan Kalai, Yanyang Ge, Mary Molacavage, and Özkan laboratory members Natalie Tsang, Svitlana Usatyuk, Wioletta Nawrocka, Indya Weathers for technical support and valuable discussions; as well as Kang Shen, Tom Clandinin, Xiaojing Gao, Judith Frydman, Xiaochen Xiong for feedback on the project and manuscript. CL was supported by the Stanford Science Fellows Program. LL is an investigator of Howard Hughes Medical Institute. This work was supported by National Institutes of Health grants R01-DC005982 (LL) and R01-NS139060 (EO).

## Author contributions

ZL, CL, and LL conceived the project. ZL, CL, and LL planed the experiments and interpreted the results. ZL, CL, CX and YH performed the *in vivo* experiments. DJL assisted in the generation of transgenic flies. ABL, JMP, and EO performed biochemical experiments. ZL, CL, and LL wrote the paper, with inputs from all other co-authors. LL supervised the work.

## Competing interests

The authors declare that they have no competing interests.

---

**Extended Data Fig. 1.**
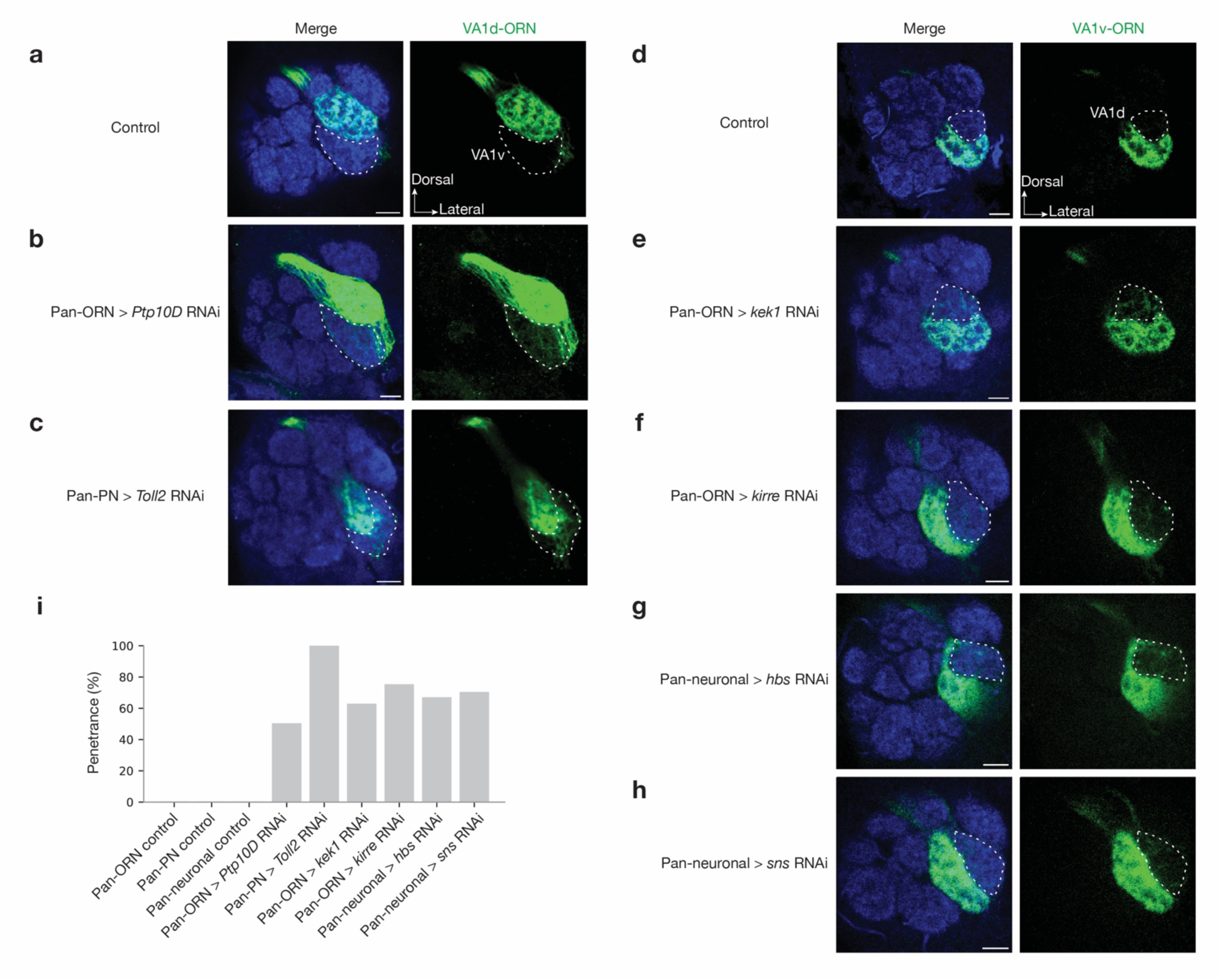
*In vivo* RNAi screen to identify CSPs required for synaptic partner matching. **a–c,** Confocal images of adult antennal lobes showing neuropil staining by N-cadherin antibody (blue) and VA1d-ORN axons (green). The VA1v glomerulus is outlined based on N-cadherin staining. Control VA1d-ORN axons only innervate the VA1d glomerulus dorsal to the VA1v glomerulus (**a**). Some VA1d-ORN axons mistarget to the VA1v glomerulus in *Ptp10D* RNAi expressed from all ORNs (**b**, maximum projection of 3 sections with 1-µm interval), or *Toll2* RNAi expressed in all PNs (**c**). **d–h,** Confocal images of adult antennal lobes showing neuropil staining by N-cadherin antibody (blue) and VA1v-ORN axons (green). The VA1d glomerulus is outlined based on N-cadherin staining. Control VA1v-ORN axons only innervate the VA1v glomerulus ventral to the VA1d glomerulus (**d**). Some VA1v-ORN axons mistarget to the VA1d glomerulus in *kek1* RNAi expressed in all ORNs (**e**), *kirre* RNAi expressed in all ORNs (**f**), *hbs* RNAi expressed in all neurons (**g**), or *sns* RNAi expressed in all neurons (**h**). **i,** Penetrance of the mistargeting phenotypes in **a–h**. For all genotypes, n ≥ 7. Scale bars = 10 µm.

**Extended Data Fig. 2.**
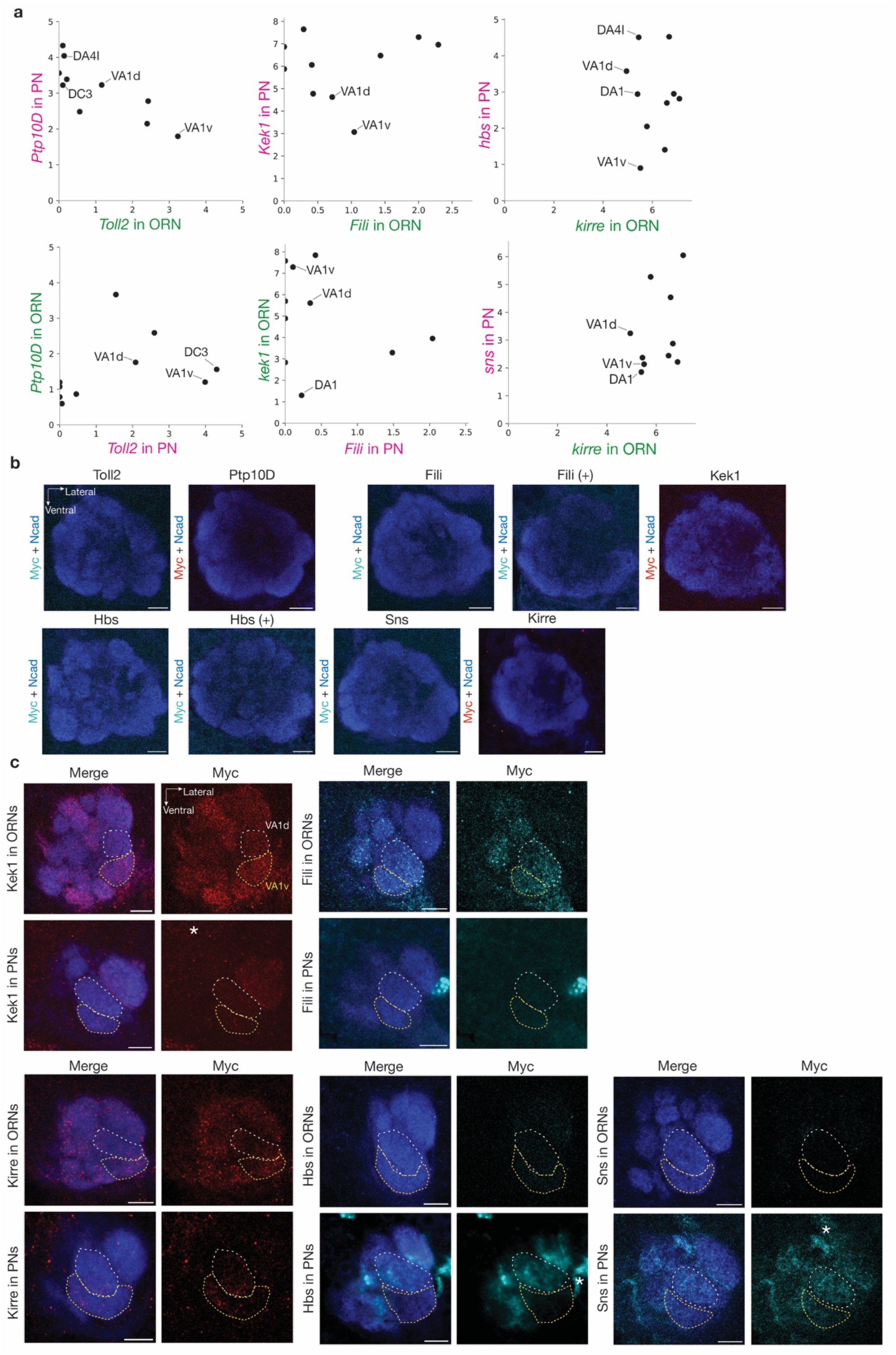
Cell-type-specific expression levels of all CSP pairs across the antennal lobe. **a,** mRNA expression levels of the three CSP pairs at 24–30h APF in ORNs and PNs that target 10 glomeruli (DA1, DA4l, DL1, DL3, DC3, VA1d, VA1v, DM2, VM3, and VA2) from single-cell RNA sequencing data^3,4^. The unit for x and y axis is log_2_(CPM+1). CPM: counts per million reads. Besides VA1d and VA1v, several other glomeruli relevant to experiments described in this study are indicated. **b**, Confocal images showing neuropil staining by N-cadherin antibody (blue) and lack of Myc staining of tagged endogenous CSPs (cyan for Toll2, Fili, Hbs, and Sns; red for Ptp10D, Kek1, and Kirre) without FLP. Annotation (+) indicate signal amplification for Myc staining. **c,** Confocal images showing neuropil staining by N-cadherin antibody (blue) and Myc staining of tagged endogenous CSPs (red for Kek1 and Kirre; cyan for Fili, Hbs, and Sns) using ORN-specific FLP (top tow) or PN-specific FLP (bottom row). The VA1d (white) and VA1v (yellow) glomeruli are outlined based on N-cadherin staining. Samples were chosen based on (1) their VA1d vs. VA1v preference indexes are close to the average, and (2) both VA1d and VA1v glomeruli occupy similar sizes in these single optic sections. Data from the above panels, along with data from Fig. 1c, d, contribute to the quantification of expression (Fig. 1e–g) and simplified schematic summary (Fig. 1h–j). Note that although Fili’s expression in PNs was undetectable using the conditional tag, a previous study showed that Fili exhibits higher expression in VA1d-PNs than VA1v-PNs using immunostaining of Fili antibody and cell-type-specific expression pattern using intersection of ORN-or PN-FLP and *Fili-GAL4*^27^. And in DA1 glomerulus, Fili appears to be expressed in a small portion of DA1-PN dendrites neighboring the VA1d glomerulus^27^. For Kirre–Hbs/Sns, expression of Hbs and Sns in ORNs are not detectable, so we did not draw their expression in Fig. 1j. We did not draw the Kirre expression level in VA1d-or VA1v-PNs (Fig. 1j) given its preference index is highly variable (Fig. 1g). We also note that the differential expression patterns of mRNAs and proteins are largely consistent (Fig. 1e–g). Occasional discrepancies could be caused by (1) post-transcriptional regulations (e.g., protein translation, stability) and (2) different time windows from which mRNA (24–30h APF) and protein (42–48h APF) data were collected. Ideally, protein staining should be done around 30h APF when synaptic partner matching initiates. However, as glomeruli have not formed at that stage, we could not distinguish cell types in which proteins are expressed. 42–48h is the earliest window we could use glomerular identity to infer cell-type-specific expression. It is possible that the expression of some of the CSPs for synaptic partner matching is already downregulated by then. Although protein expression data is more directly relevant to the action of these genes, mRNA expression data is more temporally relevant, and thus these data provide complementary information.

**Extended Data Fig. 3.**
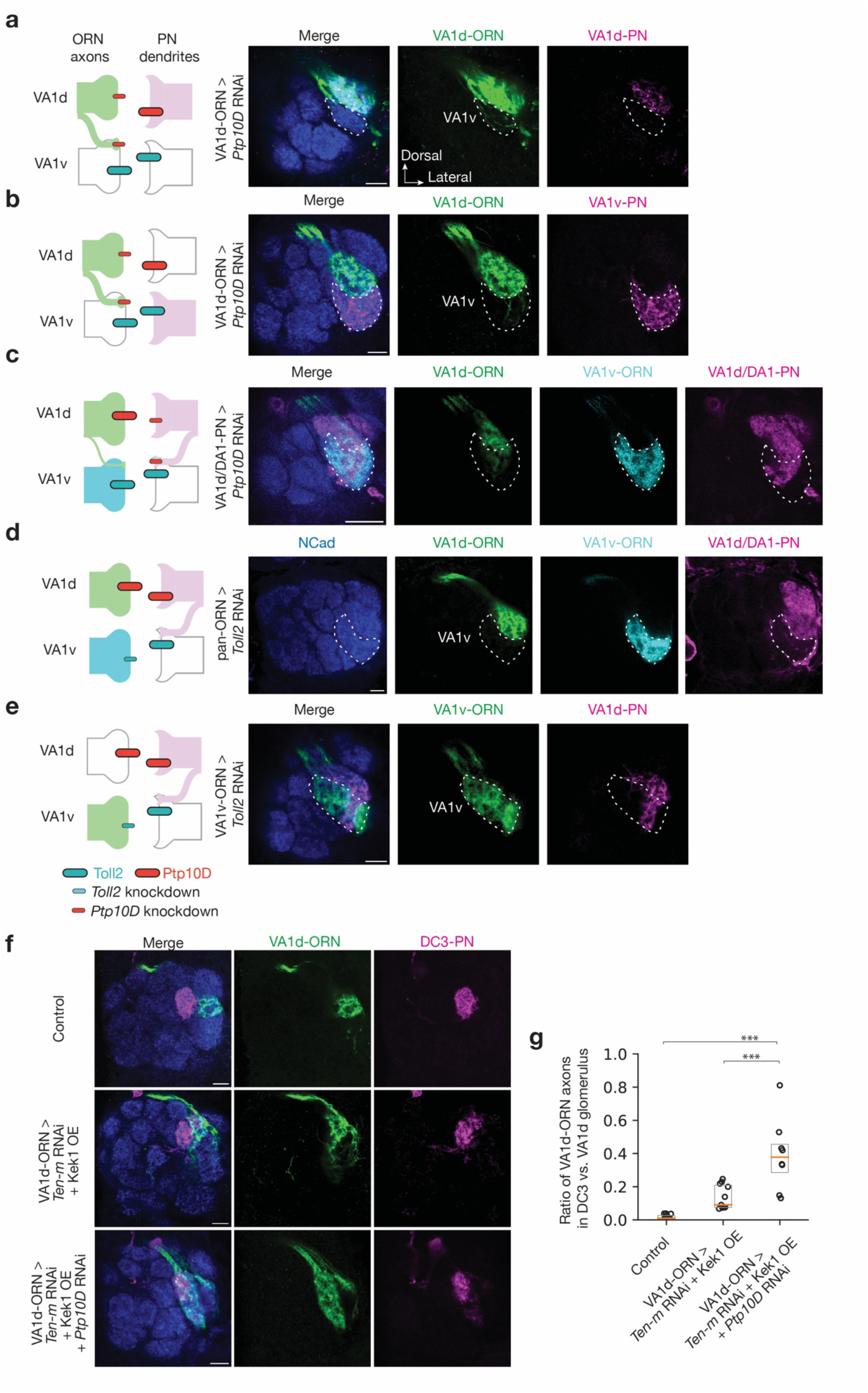
Additional loss-of-function experiments supporting that Ptp10D and Toll2 mediate PN-ORN repulsion. Schematics on the left column show genetic labeling (color-filled neurites) and genetic manipulations (red bar, high expression level of Ptp10D; cyan bar, high expression level of Toll2; smaller bar, knockdown). Right columns are confocal images of representative single sections of adult antennal lobes showing neuropil staining by N-cadherin antibody (blue), and different types of ORN axons (green or cyan) or PN dendrites (magenta). Specific glomeruli are outlined based on N-cadherin staining. **a,** To examine whether the partner neurons of VA1d-ORNs were affected in experiments described in Fig. 2d, we simultaneously labeled VA1d-PN dendrites. Although VA1d-ORN axons (green) mistarget to the VA1v glomerulus in *Ptp10D* RNAi expressed in VA1d-ORNs as in Fig. 2d, the simultaneously dual-labeled VA1d-PN dendrites (magenta) only innervate the VA1d glomerulus. This result argues against the possibility that VA1d-ORN axons mistargeting is a secondary effect of VA1d-PN dendrites mistargeting. **b,** To confirm whether mistargeted VA1d-ORNs in Fig. 2d overlap with non-partner neurons innervating VA1v glomerulus, we simultaneously labeled VA1v-PN dendrites. Mistargeted VA1d-ORN axons (green) match with the simultaneously dual-labeled VA1v-PN dendrites (magenta) in *Ptp10D* RNAi expressed in VA1d-ORNs. **c,** To examine whether VA1d-PNs’ partner and non-partner neurons were affected in Fig. 2i, we simultaneously labeled VA1d-ORN and VA1v-ORN axons. Some VA1d/DA1-PN dendrites (magenta) mistarget to the VA1v glomerulus and mismatch with VA1v-ORN axons (cyan) in *Ptp10D* RNAi expressed in VA1d/DA1-PNs as in Fig. 2i. A smaller fraction of VA1d-ORN axons (green) also mistarget to VA1v glomerulus but still intermingle with VA1d/DA1-PN dendrites. For the brains with no VA1d/DA1-PN dendrites mistargeting, no VA1d-ORN axons mistargeting was observed. This result suggests that VA1d-ORN axons mistargeting is likely to be a secondary effect of VA1d-PN dendrites mistargeting. **d,** To examine whether ORN knockdown of *Toll2* affects the correct targeting of any ORNs besides VA1d/DA1-PNs shown in Fig. 2j, we simultaneously labeled VA1d-ORN and VA1v-ORN axons with distinct markers. Whereas pan-ORN knockdown of *Toll2* causes some VA1d/DA1-PN dendrites (magenta) to mistarget to the VA1v glomerulus and mismatch with VA1v-ORN axons (cyan), consistent with VA1v-ORN knockdown of *Toll2* shown in Fig. 2j, VA1v-ORN axons only innervate the VA1v glomerulus, and VA1d-ORN axons (green) only innervate the VA1d glomerulus. This result suggests that Toll2 functions non-cell-autonomously and does not cause repulsion between VA1d-ORN axons and VA1v-ORN axons. **e,** Similar to **d**, but with a more specific manipulation as in Fig. 2j, except that we also simultaneously labeled VA1v-ORNs. Some VA1d-PN dendrites (magenta) mistarget to the VA1v glomerulus and mismatch with VA1v-ORN axons (green) when we knocked down *Toll2* only in VA1v-ORNs. This result suggests that Toll2 functions non-cell-autonomously in VA1v-ORNs to prevent VA1d-PN dendrites to mistarget to the VA1v glomerulus. **f,** To test the possibility that Ptp10D mediates homophilic attraction, we focus on DC3-PNs where Toll2 and Ptp10D are both highly expressed (Extended Data Fig. 2a). We would expect that knocking down *Ptp10D* in a potential partner ORNs would decrease its matching with DC3-PNs if Ptp10D homophilic attraction plays a more dominant role in synaptic partner matching, but would increase its matching with DC3-PNs if Ptp10D–Toll2 repulsion plays a more dominant role. We used genetic manipulations of VA1d-ORNs to test this. Confocal images show neuropil staining by N-cadherin antibody (blue), DC3-PN dendrites (magenta), and VA1d-ORN axons (green). (Top row) In control, VA1d-ORN axons only innervate the VA1d glomerulus and does not overlap with DC3-PN dendrites. (Middle row) As knocking down *Ptp10D* alone cause VA1d-ORN axons to mistarget to the VA1v glomerulus and not to DC3 glomerulus, we incorporated manipulation of Ten-m and Kek1 to sensitize VA1d-ORN based on the results in the companion manuscript^37^ Fig. 5 CSP set2. VA1d-ORN axons overlap with DC3-PN dendrites following *Ten-m* knockdown and Kek1 overexpression in VA1d-ORNs. (Bottom row) VA1d-ORN axons overlap more with DC3-PN dendrites when *Ptp10D* is knocked down. This result argues against Ptp10D mediating homophilic attraction. **g,** Quantification of the mistargeting ratio of VA1d-ORN axons overlapping with DC3-PN dendrites versus not overlapping for experiments in **f**. For all genotypes, n ≥ 10. Boxes indicate geometric mean and 25% to 75% range. Kruskal–Wallis test with Bonferroni’s multiple comparison.

**Extended Data Fig. 4.**
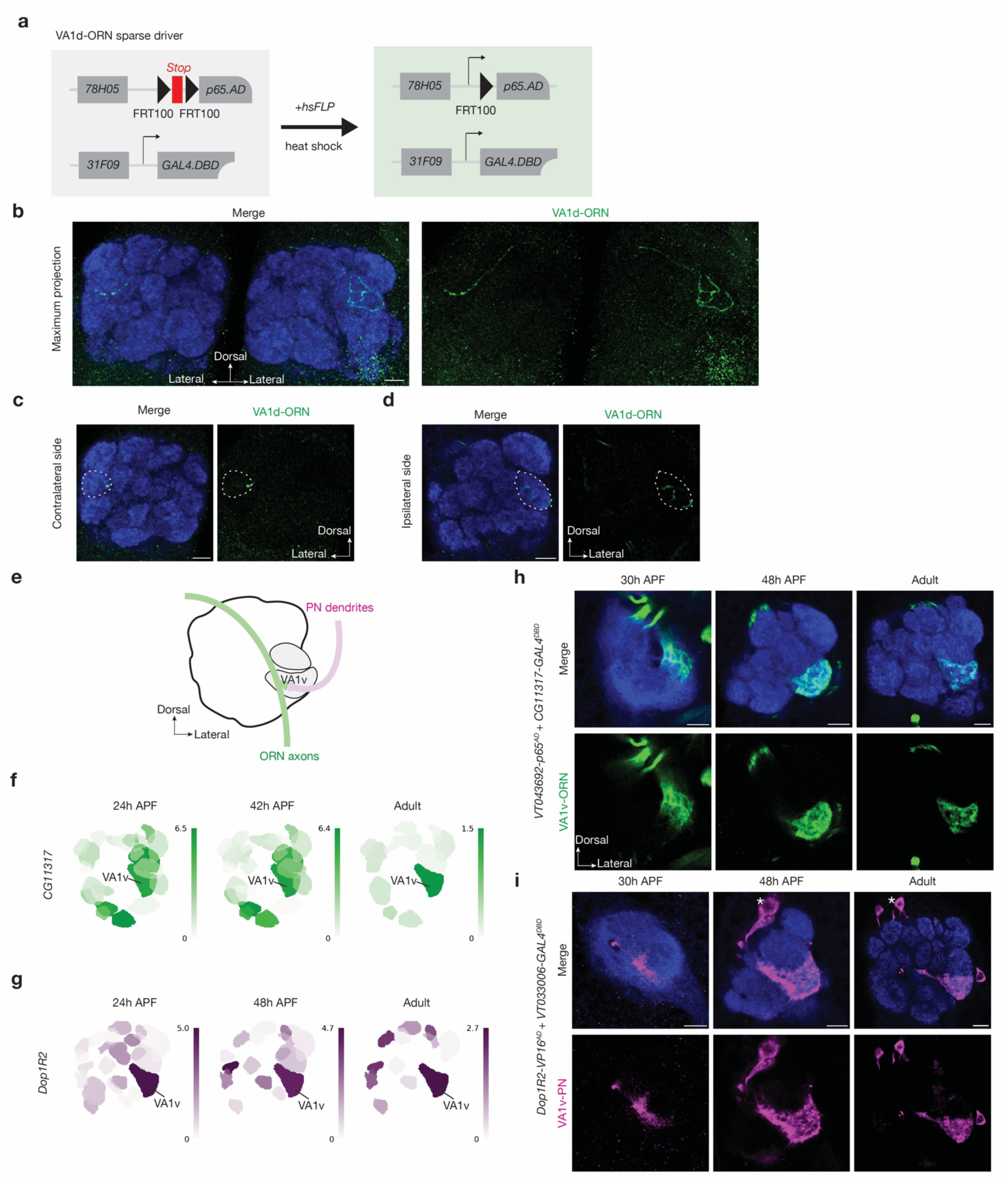
Characterization of new genetic drivers. **a,** Schematic for VA1d-ORN sparse drivers using a previously described method^67^. The transcription activation domain (AD) is controlled by the enhancer *GMR78H05* and gated by *FRT100-STOP-FRT100. FRT100* sites are ∼1% efficiency as of wild-type FRT sites. STOP represent a transcription termination sequence. Heat-shock-induced FLP expression often enables split GAL4 expression in a single VA1d-ORN. **b–d,** VA1d-ORN sparse driver enable single VA1d-ORN labeling (green, labeled by a membrane-targeted GFP) as shown in the maximum projection (**b**), single contralateral section (**c**), or single ipsilateral section (**d**). The VA1d glomerulus location is verified by neuropil staining of N-cadherin (NCad) antibody (blue). **e,** Schematic of VA1v-ORN axons and VA1v-PN dendrites matching in the VA1v glomerulus (grey) in adult *Drosophila* antennal lobe (black solid line). Green, VA1v-ORNs; Magenta, VA1v-PNs. **f, g,** Single-cell RNA sequencing data showing the expression level of *CG11317* in VA1v-ORNs (**f**) and *Dop1R2* in VA1v-PNs (**g**) are consistently high across developmental stages. Heat map units: log_2_(CPM+1). CPM: counts per million reads. Data adapted from previous published studies^3,4^. **h, i,** Gene-based genetic drivers^73^ express in VA1v-ORNs (green, labeled by a membrane-targeted GFP) (**h**) or VA1v-PNs (magenta, labeled by a membrane-targeted GFP) (**i**) at 30h APF (left), 48h APF (middle) and in adults (right). The VA1v glomerulus location is verified by neuropil staining of N-cadherin (NCad) antibody (blue). Both drivers enable expression of the transgenes in VA1v-ORNs or VA1v-PNs across developmental stages. Although they also express in ORNs or PNs targeting several other glomeruli, the expression is not detectable in glomeruli adjacent to VA1v (especially VA1d). *, PN cell bodies. Scale bars = 10 µm.

**Extended Data Fig. 5.**
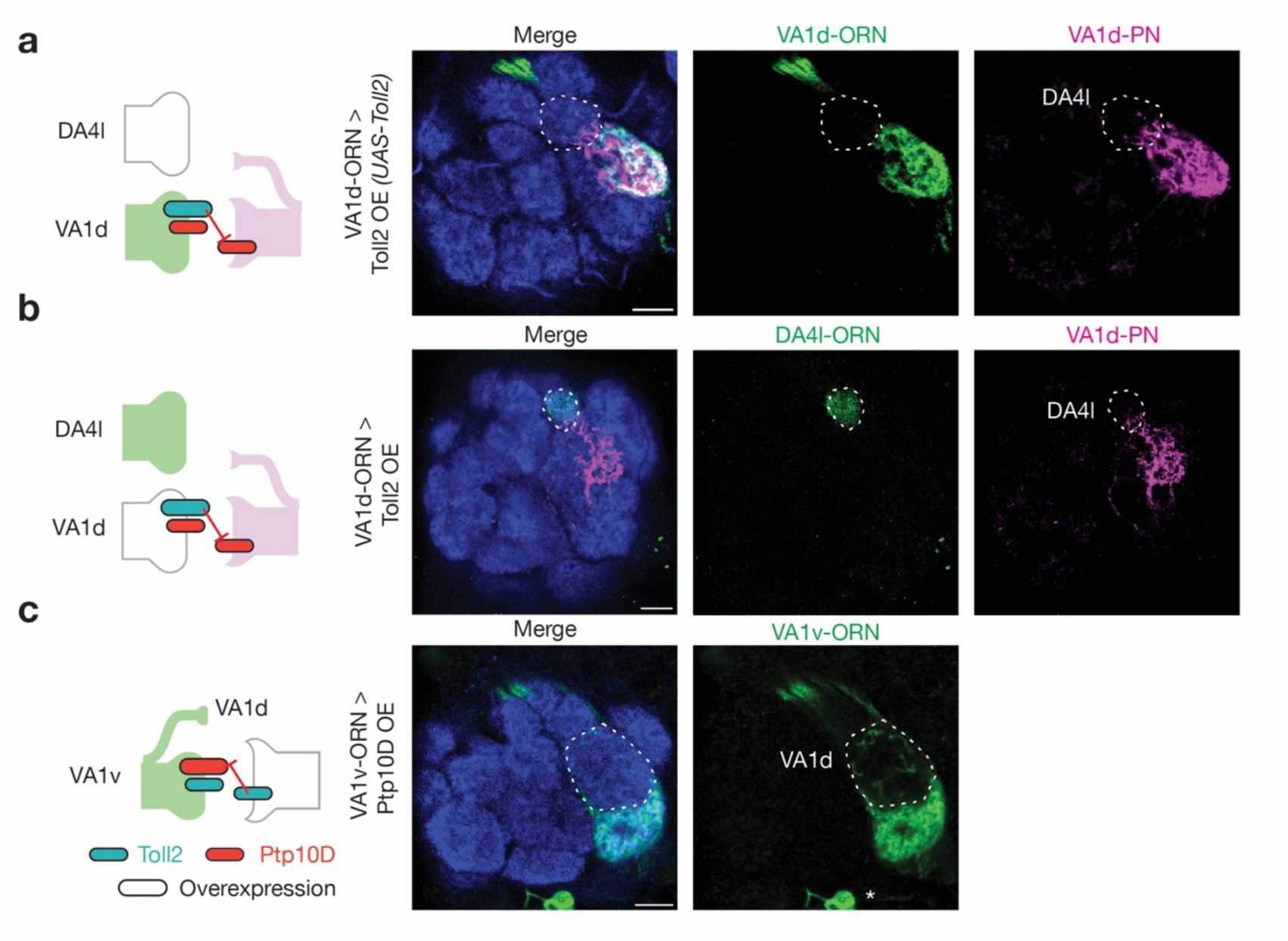
Additional gain-of-function experiments supporting that Ptp10D and Toll2 mediate PN-ORN repulsion. Schematics on the left column show genetic labeling (color-filled neurites) and genetic manipulations (red bar, high expression level of Ptp10D; cyan bar, high expression level of Toll2; large bar, overexpression). Right columns are confocal images of representative single sections of adult antennal lobes showing neuropil staining by N-cadherin antibody (blue), and different types of ORN axons (green) or PN dendrites (magenta). Specific glomeruli are outlined based on N-cadherin staining. **a,** To examine whether the partner neurons of VA1d-ORNs were affected in the experiment described in Fig. 3b, we simultaneously labeled VA1d-ORN axons. Although VA1d-PN dendrites (magenta) mistarget to the DA4l glomerulus following Toll2 overexpression in VA1d-ORNs as in Fig. 3b, VA1d-ORN axons (green) only innervate the VA1d glomerulus. This result supports a non-cell-autonomous function for Toll2. **b,** To confirm whether mistargeted VA1d-ORNs in Fig. 3b overlap with DA4l-PN dendrites in DA4l glomerulus, we simultaneously labeled DA4l-PN dendrites. Some VA1d-PN dendrites (magenta) mistarget to the DA4l glomerulus and mismatch with DA4l-ORN labeled by *Or43a-mCD8GFP* (green) following Toll2 overexpression in VA1d-ORNs. **c,** As Ptp10D expression in VA1v-ORNs is low and Toll2 expression in VA1v-PNs is high, we overexpressed Ptp10D in VA1v-ORNs, and observed that some VA1v-ORN axons (green) mistarget to the VA1d glomerulus, where both VA1d-PNs and VA1d-ORNs have low Toll2 expression (Fig. 1h). This result supports the repulsion model. Scale bars = 10 µm.

**Extended Data Fig. 6.**
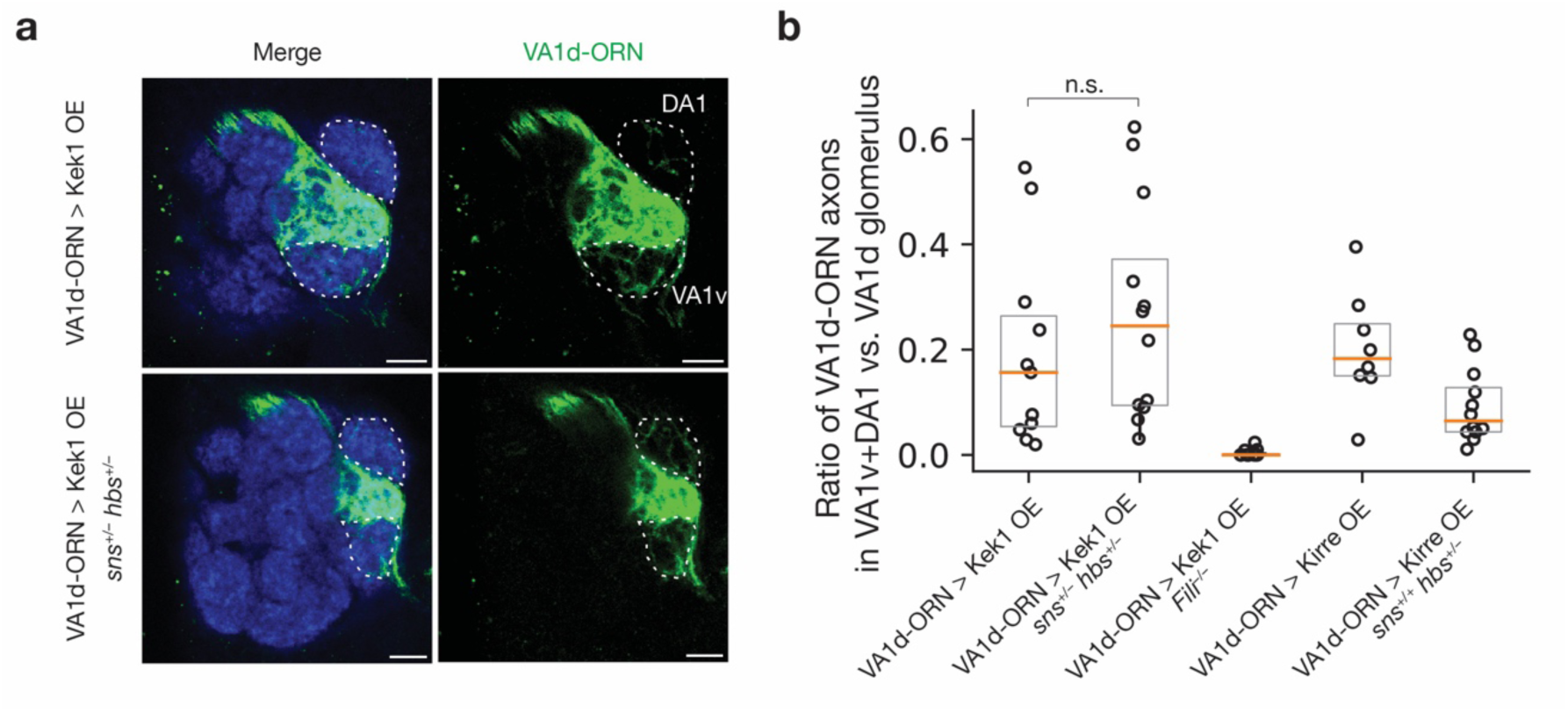
Knocking down *hbs* or *sns* does not suppress the Kek1’s gain-of-function phenotype. **a**, DA1 and VA1v glomeruli are outlined based on N-cadherin staining. Control VA1d-ORN axons only innervate the VA1d glomerulus ventral to the DA1 glomerulus and dorsal to the VA1v glomerulus (Fig. 4f). Some VA1d-ORN axons mistarget to the DA1 and VA1v glomeruli following Kek1 overexpression in VA1d-ORNs (top row). This phenotype is not suppressed in *hbs* and *sns* double heterozygous mutant (bottom row). **b,** Quantification of the mistargeting ratio of VA1d-ORN axons in the DA1+VA1v versus VA1d glomerulus. The first two columns are the quantification for panel **a**. The rest are re-plotting of part of the data in Fig. 4m and Fig. 5q, showing that the Kek1 overexpression phenotype can be suppressed by *Fili* mutant (3^rd^ column), and that *hbs* heterozygous mutant can suppress Kirre overexpression phenotype (4^th^ and 5^th^ columns). For all genotypes, n ≥ 10. Boxes indicate geometric mean and 25% to 75% range. Kruskal–Wallis test with Bonferroni’s multiple comparison.

**Extended Data Fig. 7.**
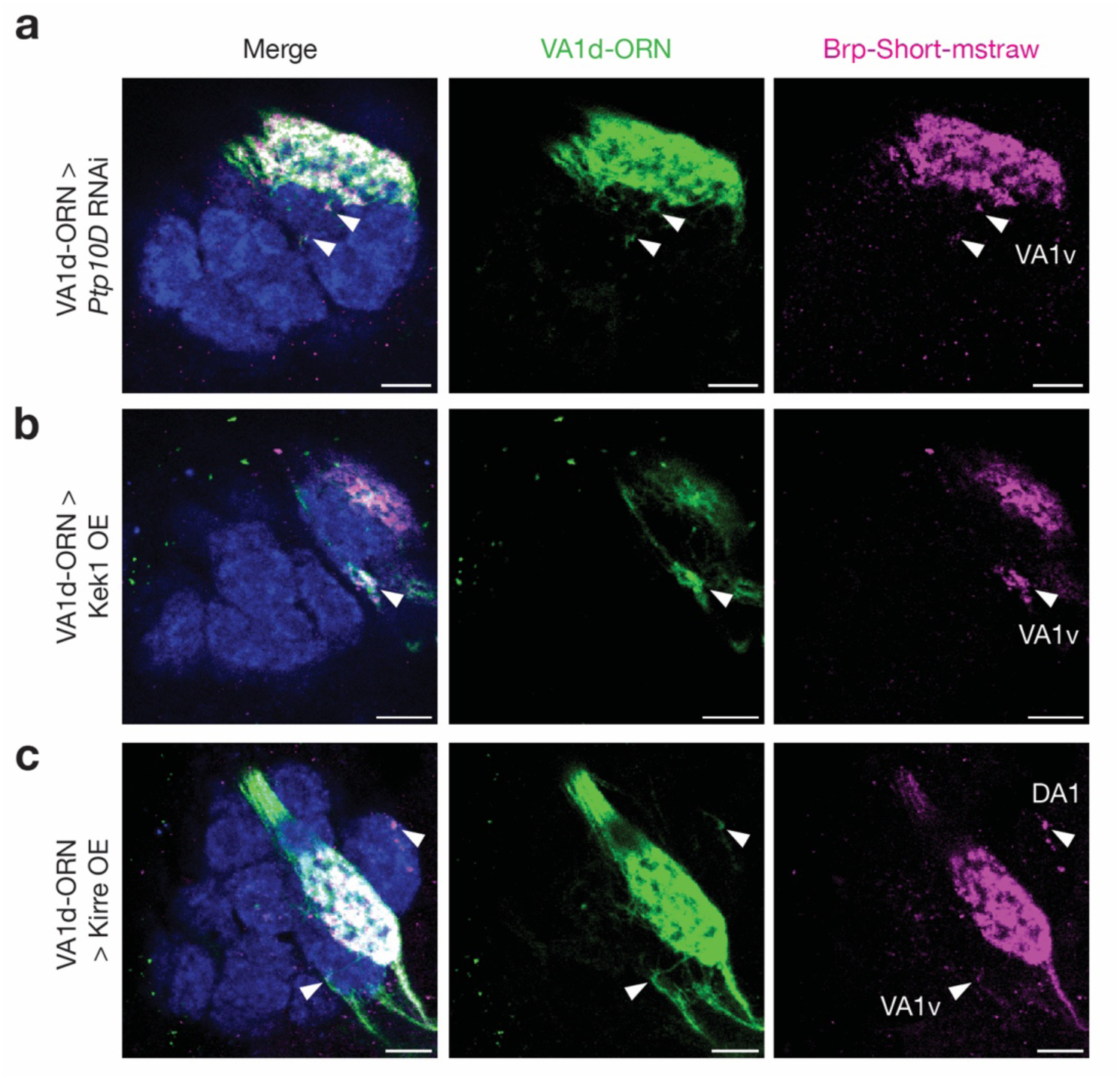
Mistargeted VA1d-ORN axons in neighboring glomeruli are enriched for a presynaptic terminal protein. Confocal images of adult antennal lobes showing neuropil staining by N-cadherin antibody (blue), VA1d-ORN axons (green, labeled by a membrane-targeted GFP) and a presynaptic active zone marker Bruchpilot-Short^74^ (Brp-Short, magenta). Knockdown of *Ptp10D* (**a**), overexpression of Kek1 (**b**), and overexpression of Kirre (**c**) in VA1d-ORNs caused their axons to mistarget to neighboring glomeruli (arrowheads). These mistargeted processes are enriched for Brp-Short, suggesting that mistargeted ORN axons might form synapses. Scale bar = 10 µm.

**Extended data Fig. 8.**
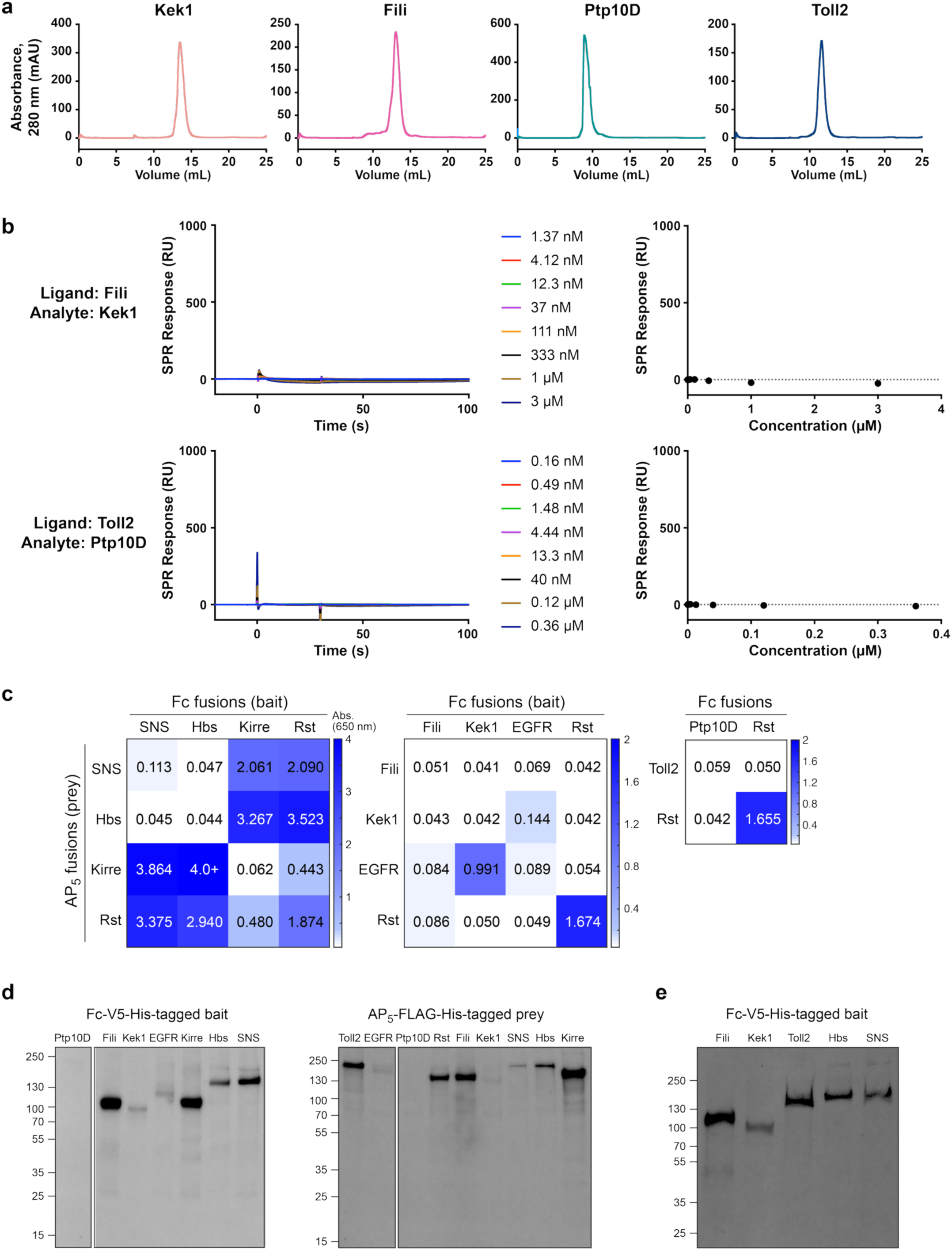
*In vitro* assays to test direct binding between CSP pairs. **a,** Size-exclusion chromatography curves for four extracellular domains (ectodomains) purified on a Superdex 200 Increase 10/300 column on an ÄKTA Pure system, reporting absorbance at 280 nm with a path length of 0.2 cm. **b,** Surface plasmon resonance sensorgams of 30 s injections (left) and analyte concentration vs. response plots (right) show no direct physical interaction between Fili and Kek1, and between Toll2 and Ptp10D. **c,** Extracellular interactome assay (ECIA) to test binding between various ectodomains studied. We observe strong binding between Sns and Hbs with Kirre and its paralog Rst (left), and Kek1 with its previously described binding partner, EGFR (middle). No interaction between Fili and Kek1 (middle) or Ptp10D and Toll2 (right) were observed. **d,** Western blots of proteins used in the ECIA in (**c**) against the hexahistidine tag common in all constructs. There is a lack of detectable expression for Ptp10D ectodomain, which may be the reason behind no binding for Ptp10D in (**c**). **e,** Western blots of the same Fc fusions used directly in the tissue staining experiments in Extended Data Fig. 9.

**Extended data Fig. 9.**
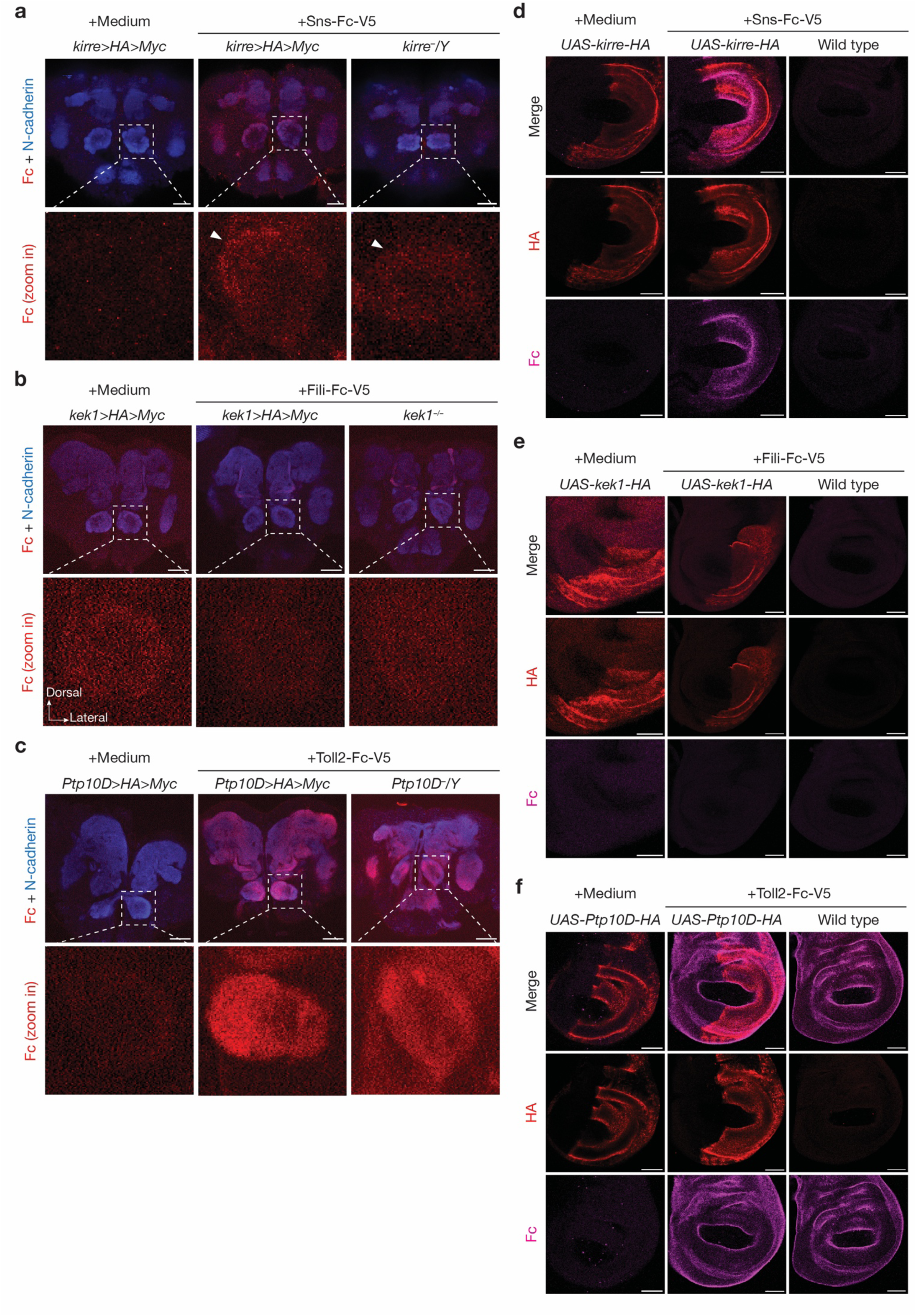
Live tissue staining assays to test direct binding between CSP pairs. **a,** Confocal images of pupal brains (48h APF) showing neuropil staining by N-cadherin antibody (blue), and binding of Sns-Fc-V5 in live pupa brains followed by fixing and staining against Fc (red). Control *kirre* conditionally tagged brains incubated with medium alone showed minimal background signal for Fc staining (left column). *kirre* conditionally tagged brains incubated with Sns-Fc-V5 proteins in medium have detectable signal for Fc staining in multiple brain regions and have differential signal in different glomeruli (middle column). These signals are substantially diminished in *kirre* mutant brains incubated with Sns-Fc-V5 proteins (right column), especially in the glomeruli where signal is high in the *kirre* conditionally tagged brains (arrowheads in the magnified images in the bottom rows). These results suggest Sns-Fc-V5 binds to endogenous Kirre, which serve as a positive control for the binding assay. **b,** Confocal images of pupal brains (48h APF) showing neuropil staining by N-cadherin antibody (blue), and binding of Fili-Fc-V5 in live pupal brains followed by fixing and staining against Fc (red). *kek1* conditionally tagged brains (middle column) incubated with Fili-Fc-V5 proteins in medium do not exhibit an increase of Fc signal in the antennal lobe (magnified in the bottom row) comparing to the control without the addition of Fili-Fc-V5 (left column) or Fili-Fc-V5 in *kek1* homozygous mutant brain (right column). These results suggest that Kek1 does not bind to Fili in this assay. **c,** Confocal images of pupal brains (48h APF) showing neuropil staining by N-cadherin antibody (blue), and binding of Toll2-Fc-V5 in live pupa brains followed by fixing and staining against Fc (red). Control *Ptp10D* conditionally tagged brains incubated with medium alone showed minimal background signal for Fc staining (left column). *Ptp10D* conditionally tagged brains incubated with Toll2-Fc-V5 proteins in medium have detectable signal for Fc staining in multiple brain regions (middle column). However, these signals are still present in *Ptp10D* mutant brains incubated with Toll2-Fc-V5 proteins (right column). These results suggest Toll2-Fc-V5 binds to other proteins expressed in the brain, masking the detection of its potential binding to Ptp10D. **d**, Confocal images of wing disc dissected from 3rd larvae showing ectopic expression of *UAS-kirre-HA* in the posterior compartment driven by *engrailed (en)-GAL4* (HA staining, red), and binding of Sns-Fc-V5 in live larvae followed by fixing and staining against Fc (magenta). Control Kirre-overexpressed wing disc incubated with medium alone shows minimal background signal for Fc staining (left column). Kirre-overexpressed wing disc incubated with Sns-Fc-V5 proteins in medium has specific binding signal in the posterior compartment where Kirre was overexpressed from *en-GAL4* (middle column). Interestingly, Sns binds strongest to regions with intermediate but not highest levels of Kirre overexpression. Without Kirre overexpression, no specific signal in the posterior compartment is detected (right column). These results suggest Sns-Fc-V5 binds to Kirre, which serves as a positive control for the binding assay. **e**, Confocal images of wing disc dissected from 3rd larvae showing ectopic expression of *UAS-kek1-HA* in the posterior compartment driven by *en-GAL4* (HA staining, red), and binding of Fili-Fc-V5 in live larvae followed by fixing and staining against Fc (magenta). Fc signal is not detectable in Kek1-overexpressed wing disc incubated with medium alone, Kek1-overexpressed wing disc incubated with Fili-Fc-V5, or wild-type wing disc incubated with Fili-Fc-V5. These results suggest that Kek1 does not bind to Fili in this assay. **f**, Confocal images of wing disc dissected from 3rd stage larvae showing ectopic expression of *UAS-Ptp10D-HA* in the posterior compartment driven by *en-GAL4* (HA staining, red), and binding of Toll2-Fc-V5 in live larvae followed by fixing and staining against Fc (magenta). Control Ptp10D-overexpressed wing disc incubated with medium alone showed minimal background signal for Fc staining (left column). Ptp10D-overexpressed wing disc incubated with Toll2-Fc-V5 proteins in medium has binding signal throughout the wing disc, not restricted to the posterior compartment where Toll2 was overexpressed (middle column). These signals are still present in wild-type wing disc incubated with Toll2-Fc-V5 proteins (right column). These results suggest Toll2-Fc-V5 binds to other proteins expressed in the wing disc, masking the detection of its potential binding to Ptp10D. The Fc-V5-tagged proteins used in this figure are visualized on Western blots in Extended Data Fig. 8e. Scale bar = 20 µm (**a–c**); Scale bar = 50 µm (**d–f**).

**Extended Data Fig. 10.**
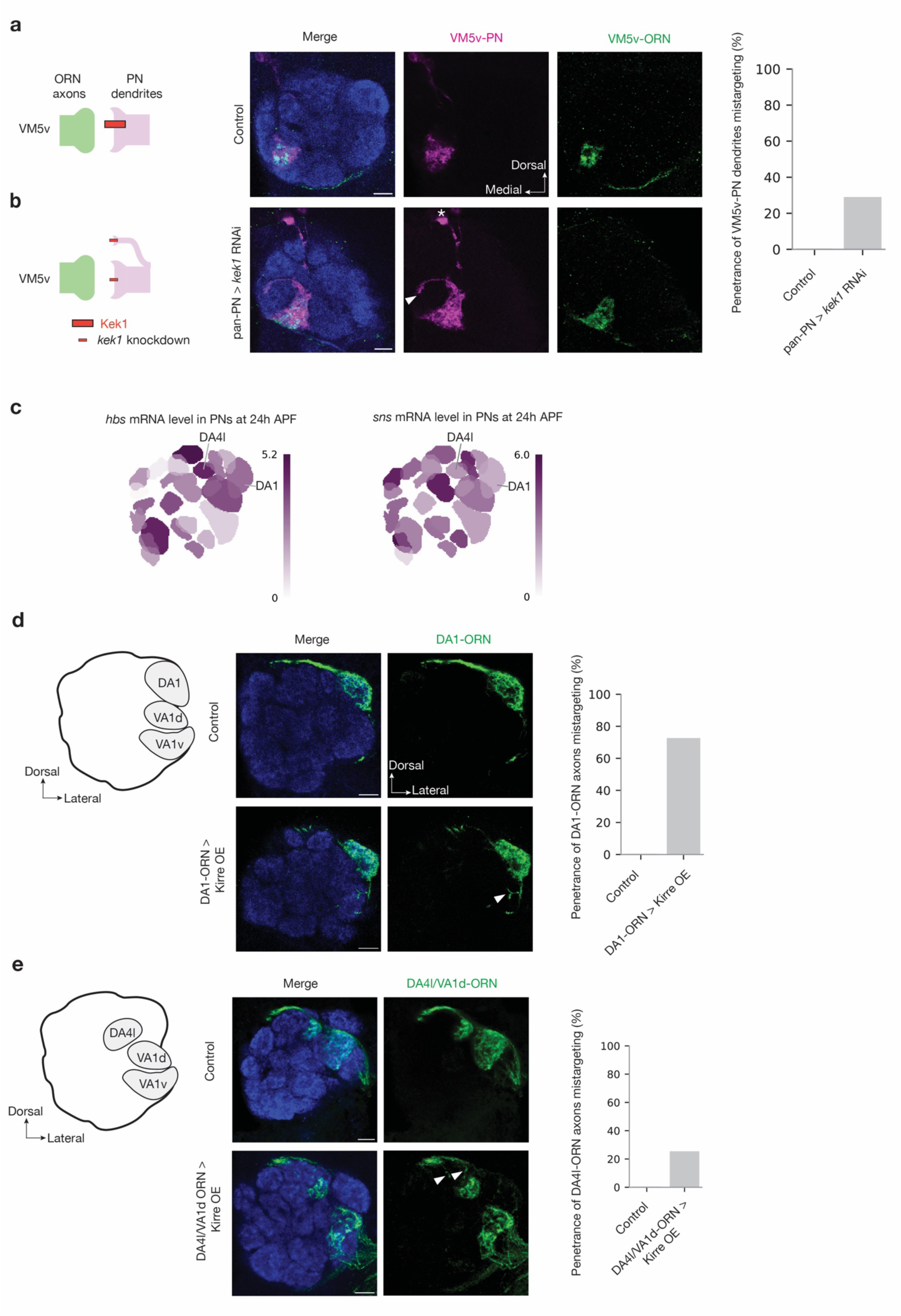
Genetic manipulation experiments of the CSPs in other parts of the antennal lobe. a, b, Left column shows experimental schematic (red bar, high expression level of Kek1; smaller bar, knockdown). The middle columns are confocal images showing adult antennal lobe neuropil staining by N-cadherin antibody (blue), VM5v-PN dendrites (magenta), and VM5v-ORN axons (green). Control VM5v-PN dendrites labeled by *GMR86C10-LexA* only innervate the VM5v glomerulus and fully overlap with VM5v-ORN axons labeled by *Or98a-mCD8GFP* (**a**). Some VA5v-PN dendrites mistarget to other glomerulus (arrowheads) in *kek1* RNAi driven by the pan-PN driver, phenocopying ORN knockdown of *Fili*^27^ (**b**). * denotes PN cell body. The right column shows penetrance of the mistargeting phenotypes. **c,** Single-cell RNA sequencing data showing the expression level of *hbs* and *sns* throughout the adult antennal lobe at 24–30 h APF. Heat map units: log_2_(CPM+1). CPM: counts per million reads. Data adapted from previous work^3,4^. **d, e,** Left column shows schematic of the adult antennal lobe with locations of three glomeruli highlighted in grey. Middle columns are confocal images showing adult antennal lobe neuropil staining by N-cadherin antibody (blue), DA1-ORN axons (green in **d**), and DA4l/VA1d-ORN axons (green in **e**). Some DA1-ORN axons or DA4l/VA1d-ORN axons mistarget to neighboring glomeruli (bottom rows) following Kirre overexpression (arrowheads), which is not observed in control (top rows). Right column shows penetrance of mistargeting phenotypes. For all genotypes, n ≥ 10. Scale bars = 10 µm.

**Extended Data Table 1.** Information for genes and manipulations used in the genetic screen.

| Gene | RNAi source | RNAi code | Manipulation | Penetrance | Gene | RNAi source | RNAi code | Manipulation | Penetrance |
| --- | --- | --- | --- | --- | --- | --- | --- | --- | --- |
| *CCKLR-17D3* | BDSC | 67866 | Mz19-PN | 0/6 | *klg* | BDSC | 28746 | Mz19-PN | 0/7 |
| *CCKLR-17D3* | BDSC | 67866 | VA1d-ORN | 4/5 | *klg* | BDSC | 28746 | pan-PN | 0/7 |
| *CG10474* | BDSC | 51444 | pan-PN | 0/7 | *klg* | BDSC | 28746 | pan-ORN | 6/9* |
| *CG14024* | VDRC | 100991 | pan-ORN | 0/5* | *klg* | VDRC | 102502 | pan-ORN | 0/6* |
| *CG33143* | BDSC | 28823 | pan-PN | 0/12* | *klg* | VDRC | 108818 | pan-ORN | 0/7* |
| *Con* | VDRC | 17898 | pan-ORN | 7/7 | *Mcr* | BDSC | 65896 | pan-ORN | 0/3* |
| *Con* | BDSC | 28967 | pan-ORN | 4/5* | *Mcr* | BDSC | 65896 | pan-PN | 3/4* |
| *Con* | BDSC | 28967 | pan-PN | 3/7 | *mgl* | BDSC | 29324 | pan-ORN | 3/3* |
| *Cow* | BDSC | 38235 | pan-ORN | 1/7* | *mgl* | VDRC | 36389 | pan-ORN | 0/7* |
| *Cow* | BDSC | 44530 | pan-ORN | 0/9* | *Nrt* | VDRC | 8495 | pan-ORN | 0/8* |
| *Cow* | BDSC | 44530 | pan-PN | 1/8* | *Nrt* | BDSC | 28742 | pan-ORN | 0/10* |
| *cue* | VDRC | 1043 | pan-ORN | 5/5* | *Nrt* | VDRC | 106080 | pan-ORN | 0/6* |
| *cue* | VDRC | 104645 | pan-ORN | 0/5* | *Nrt* | BDSC | 28742 | pan-PN | 0/7* |
| *cue* | BDSC | 36875 | pan-ORN | 0/7 | *PlexB* | BDSC | 57813 | Mz19-PN | 3/8 |
| *Diap2* | VDRC | 101187 | pan-ORN | 0/6* | *PlexB* | BDSC | 57813 | VA1d-ORN | 0/5 |
| *Dop1R1* | BDSC | 62193 | pan-PN | 0/7 | *Ptp10D* | VDRC | 8010 | pan-ORN | 4/8 |
| *ed* | BDSC | 38209 | pan-ORN | 0/7* | *Ptp10D* | BDSC | 39001 | pan-ORN | 8/13* |
| *ed* | BDSC | 38243 | pan-ORN | 0/4* | *Ptp10D* | BDSC | 39001 | pan-PN | 7/7* |
| *ed* | BDSC | 38243 | pan-PN | 3/7* | *Ptp99A* | BDSC | 57299 | DA1-ORN | 7/7 |
| *Fas2* | BDSC | 34084 | Mz19-PN | 0/7* | *Ptp99A* | BDSC | 57299 | Mz19-PN | 7/7 |
| *Fas2* | BDSC | 34084 | pan-ORN | 0/10* | *Ptp99A* | BDSC | 57299 | pan-ORN | 2/10* |
| *Fas2* | VDRC | 103807 | pan-ORN | 0/7* | *Ptp99A* | VDRC | 103457 | pan-ORN | 0/8* |
| *Fas2* | BDSC | 28990 | pan-ORN | 0/8* | *Ptp99A* | VDRC | 108505 | pan-ORN | 0/7* |
| *Fas3* | BDSC | 77396 | pan-PN | 0/7 | *Ptp99A* | BDSC | 57299 | pan-PN | 3/7 |
| *Fas3* | BDSC | 77396 | pan-ORN | 0/5* | *rst\*\** | BDSC | 28672 | DA1-ORN | 0/7 |
| *fra* | BDSC | 40826 | pan-PN | 0/7 | *rst\*\** | BDSC | 28672 | Mz19-PN | 0/7 |
| *fred* | BDSC | 42621 | DA1-ORN | 0/6 | *rst\*\** | BDSC | 28672 | pan-PN | 0/7 |
| *fred* | BDSC | 42621 | Mz19-PN | 2/2 | *sdk* | BDSC | 33412 | Mz19-PN | 1/6 |
| *fred* | BDSC | 42621 | pan-ORN | 0/9* | *sdk* | BDSC | 33412 | VA1d-ORN | 0/5 |
| *fred* | VDRC | 46180 | pan-ORN | 1/6* | *side-IV* | BDSC | 82064 | VA1d-ORN | 10/10 |
| *fred* | VDRC | 101138 | pan-ORN | 0/8* | *side-IV* | VDRC | 102563 | VA1d-ORN | 5/7 |
| *fred* | BDSC | 42621 | pan-PN | 0/7* | *side-IV* | BDSC | 82064 | VA1d-PN | 1/8 |
| *hbs* | VDRC | 40898 | pan-neuronal | 8/12 | *side-IV* | VDRC | 102563 | VA1d-PN | 2/12 |
| *hbs* | BDSC | 57003 | pan-PN | 3/6* | *side-V* | BDSC | 61953 | Mz19-PN | 2/9 |
| *ldit* | VDRC | 101893 | pan-ORN | 0/6* | *side-V* | BDSC | 61953 | VA1d-ORN | 6/6 |
| *ldit* | BDSC | 28823 | pan-ORN | 0/12* | *slit* | VDRC | 20210 | pan-ORN | 0/7* |
| *ldit* | BDSC | 28823 | pan-PN | 0/4* | *slit* | BDSC | 31467 | pan-ORN | 3/10* |
| *kek1* | VDRC | 36252 | pan-ORN | 5/6* | *slit* | BDSC | 31468 | pan-ORN | 2/8* |
| *kek1* | BDSC | 57000 | pan-ORN | 5/8 | *slit* | VDRC | 108853 | pan-ORN | 0/7* |
| *kek1* | VDRC | 101166 | pan-ORN | 1/8* | *slit* | BDSC | 31467 | pan-PN | 0/7* |
| *kek5* | BDSC | 40830 | Mz19-PN | 0/7 | *slit* | BDSC | 31468 | pan-PN | 0/7* |
| *kek5* | BDSC | 40830 | pan-ORN | 7/7* | *sns* | VDRC | 109442 | pan-neuronal | 14/20 |
| *kek5* | BDSC | 40830 | pan-PN | 2/7* | *TkR86C* | VDRC | 13392 | pan-ORN | 0/5* |
| *kek5* | BDSC | 40830 | VA1d-ORN | 4/5 | *TkR86C* | VDRC | 107090 | pan-ORN | 0/6* |
| *kek6* | BDSC | 61212 | Mz19-PN | 0/7 | *TkR86C* | BDSC | 31884 | pan-ORN | 0/3* |
| *kek6* | BDSC | 61212 | pan-ORN | 7/7 | *tnc* | BDSC | 60058 | pan-PN | 7/7* |
| *kek6* | VDRC | 109681 | VA1d-ORN | 5/6 | *Toll2* | VDRC | 965 | pan-ORN | 8/8* |
| *kirre* | VDRC | 109585 | pan-ORN | 9/12 | *Toll2* | BDSC | 30498 | pan-ORN | 2/3* |
|  |  |  |  |  | *Toll2* | VDRC | 36305 | pan-ORN | 2/3* |
|  |  |  |  |  | *Toll2* | VDRC | 965 | pan-PN | 3/7 |

\* indicates simultaneous knockdown of *Ten-m* using RNAi. Some of the screening experiments were done with the simultaneous knockdown of *Ten-m* to ‘sensitize’ the system because: (1) Knocking down *Ten-m* causes partial mismatching phenotype between ORN axons and PN dendrites<sup>13</sup> and (2) *Ten-m* is highly expressed in many of the glomeruli labeled in our screening. In some of the ‘sensitized’ screening experiments, we followed up the initial phenotype with single manipulation that did not contain *Ten-m* knockdown. In general, we observed equal or higher penetrance when *Ten-m* was simultaneously knocked down. For example, in the *Toll2* RNAi experiments, the penetrance is 3/7 when using RNAi line 965 from VDRC alone and is 8/8 when using both line 965 and *Ten-m* RNAi.
\*\* Rst (Roughest) binds with Hbs and Sns *in vitro*<sup>49</sup> and *rst* is also expressed in PNs and ORNs based on the single-cell transcriptome data<sup>3,4</sup>. However, we did not observe any mistargeting phenotype in cell-type-specific *rst* knockdown experiments.

**Extended Data Table 2.** Summary of genetic manipulation experiments that did not produce mistargeting phenotypes.

| Ex p.# | Genetic manipulation | Cell types examined | m/n<sup>1</sup> | Interpretation |
| --- | --- | --- | --- | --- |
| 1 | *Ptp10D*<sup>-</sup>/*Y* | VA1v-ORN axons | 0/18 | Ptp10D is not required for VA1v-ORN axon targeting |
| 2 | VA1d/DA1-PN > *Ptp10D* RNAi | VA1d-ORN axons | 0/14 | Ptp10D does not affect targeting of partner neurons |
| 3 | VA1v-ORN > *Ptp10D* RNAi | VA1v-ORN axons | 0/10 | Ptp10D does not function in VA1v-ORNs for their axon targeting |
| 4 | VA1d/DA1-PN > *Toll2* RNAi | VA1d-ORN axons | 0/12 | Toll2 does not affect targeting of partner neurons |
| 5 | VA1d/DA1-PN > *Toll2* RNAi | VA1v-ORN axons | 0/12 | Toll2 does not function in VA1d-PNs to regulate VA1v-ORN axon targeting |
| 6 | VA1d/DA1-PN > *Toll2* RNAi | VA1d/DA1 PN dendrites | 0/12 | Toll2 does not function in VA1d-PNs to regulate their dendrite targeting |
| 7 | pan-ORN > *Toll2* RNAi | VA1v-PN dendrites | 0/10 | Toll2 does not affect targeting of partner neurons |
| 8 | VA1v-ORN > *Toll2* RNAi | VA1v-PN dendrites | 0/10 | Toll2 does not affect targeting of partner neurons |
| 9 | VA1d-ORN > *Toll2* OE | VA1v-PN dendrites | 0/11 | Toll2 unlikely mediates homophilic attraction |
| 10 | VA1d/DA1-PN > *Fili* RNAi | VA1d-ORN axons | 0/14 | Fili does not affect targeting of partner neurons |
| 11 | *kek1*<sup>-/-</sup> | VA1d-ORN axons | 0/18 | Kek1 is not required for VA1d-ORN axon targeting |
| 12 | pan-ORN > *kek1* RNAi | VA1d-ORN axons | 0/10 | Kek1 is not required in ORNs for in VA1d-ORN axon targeting |
| 13 | VA1d-ORN > *kek1* RNAi | VA1d-ORN axons | 0/10 | Kek1 is not required in VA1d-ORNs for their axon targeting |
| 14 | pan-ORN > *kek1* RNAi | VA1v-PN dendrites | 0/11 | Kek1 does not affect targeting of partner neurons |
| 15 | VA1v-ORN > *kek1* RNAi | VA1v-PN dendrites | 0/10 | Kek1 does not affect targeting of partner neurons |
| 16 | VA1d-ORN > *Kek1* OE | VA1v-PN dendrites | 0/10 | Kek1 unlikely mediates homophilic attraction |
| 17 | VA1v-ORN > *Kek1* OE | VA1v-ORN axons | 0/10 | Kek1 endogenous expression level is higher in VA1v-ORNs than VA1d-ORNs and overexpression of Kek1 in VA1v-ORNs does not cause ORN axon mistargeting phenotype |
| 18 | *hbs*<sup>+/-</sup> | VA1d-ORN axons | 0/10 | Mutant alone does not have any VA1d-ORN axons mistargeting phenotype (Control for Fig. 5) |
| 19 | *hbs*<sup>+/-</sup>, *sns*<sup>-/-</sup> | VA1d-ORN axons | 0/10 | Mutant alone does not have any VA1d-ORN axons mistargeting phenotype (Control for Fig. 5) |
| 20 | VA1d/DA1-PN > *hbs* RNAi | VA1d-ORN axons | 0/10 | Hbs does not affect targeting of partner neurons |
| 21 | *sns*<sup>-/-</sup> | VA1d-ORN axons | 0/10 | Mutant alone does not have any VA1d-ORN axons mistargeting phenotype (Control for Fig. 5) |
| 22 | VA1d/DA1-PN > *sns* RNAi | VA1d-ORN axons | 0/10 | Sns does not affect targeting of partner neurons |
| 23 | *kirre*<sup>-</sup>/Y | VA1d-ORN axons | 0/10 | Mutant alone does not have any VA1d-ORN axons mistargeting phenotype (Control for Fig. 5) |
| 24 | pan-ORN > *kirre* RNAi | VA1d-ORN axons | 0/12 | Kirre is not required in ORNs for VA1d-ORN axon targeting |
| 25 | VA1d-ORN > *kirre* RNAi | VA1d-ORN axons | 0/10 | Kirre is not required in VA1d-ORNs for their axon targeting |
| 26 | pan-ORN > *kirre* RNAi | VA1v-PN dendrites | 0/15 | Kirre does not affect targeting of partner neurons |
| 27 | VA1v-ORN > *kirre* RNAi | VA1v-PN dendrites | 0/10 | Kirre does not affect targeting of partner neurons |

<sup>1</sup> m = number of antennal lobes with mistargeting phenotypes; n = number of total antennal lobes examined.

**Extended Data Table 3.**
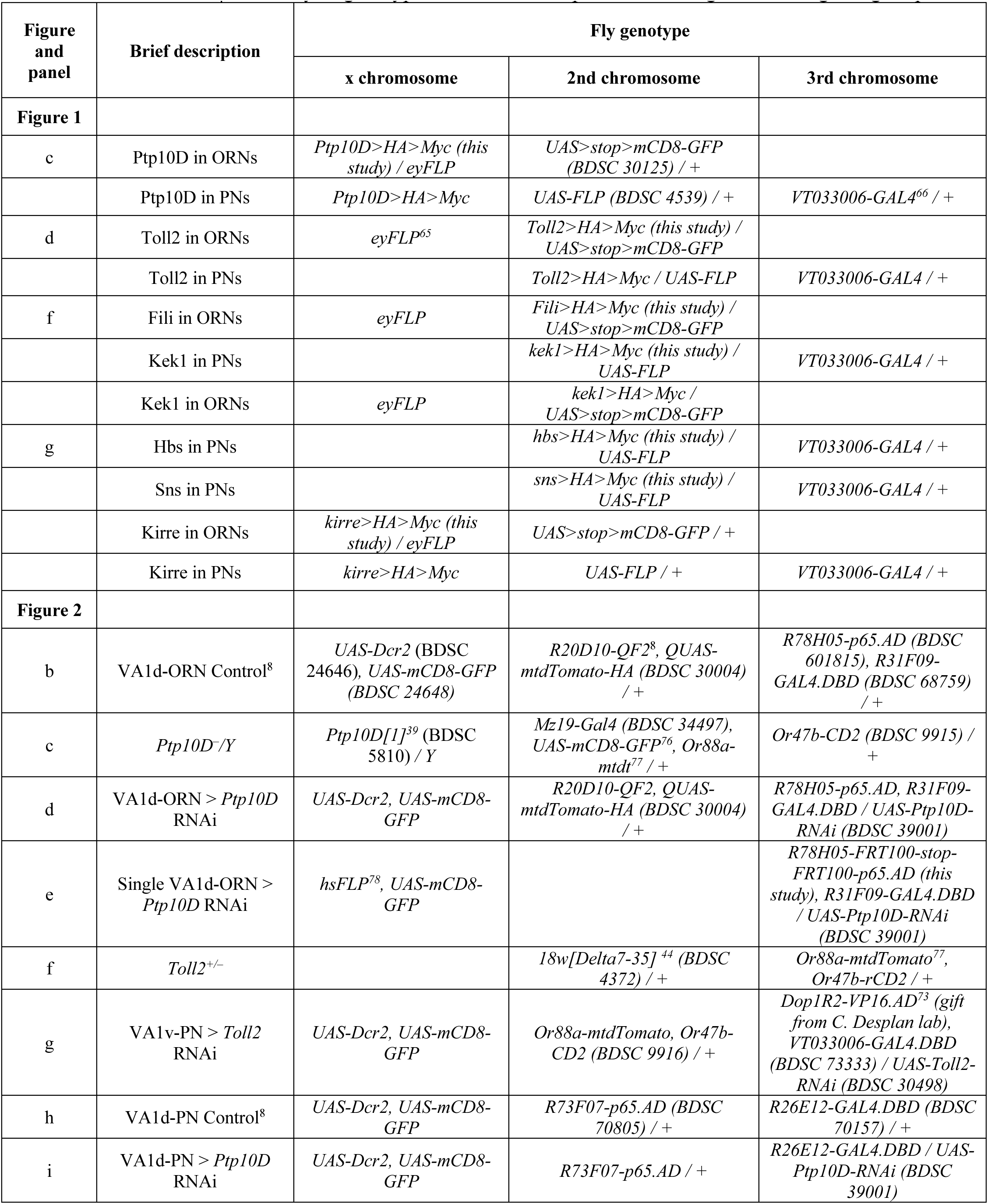
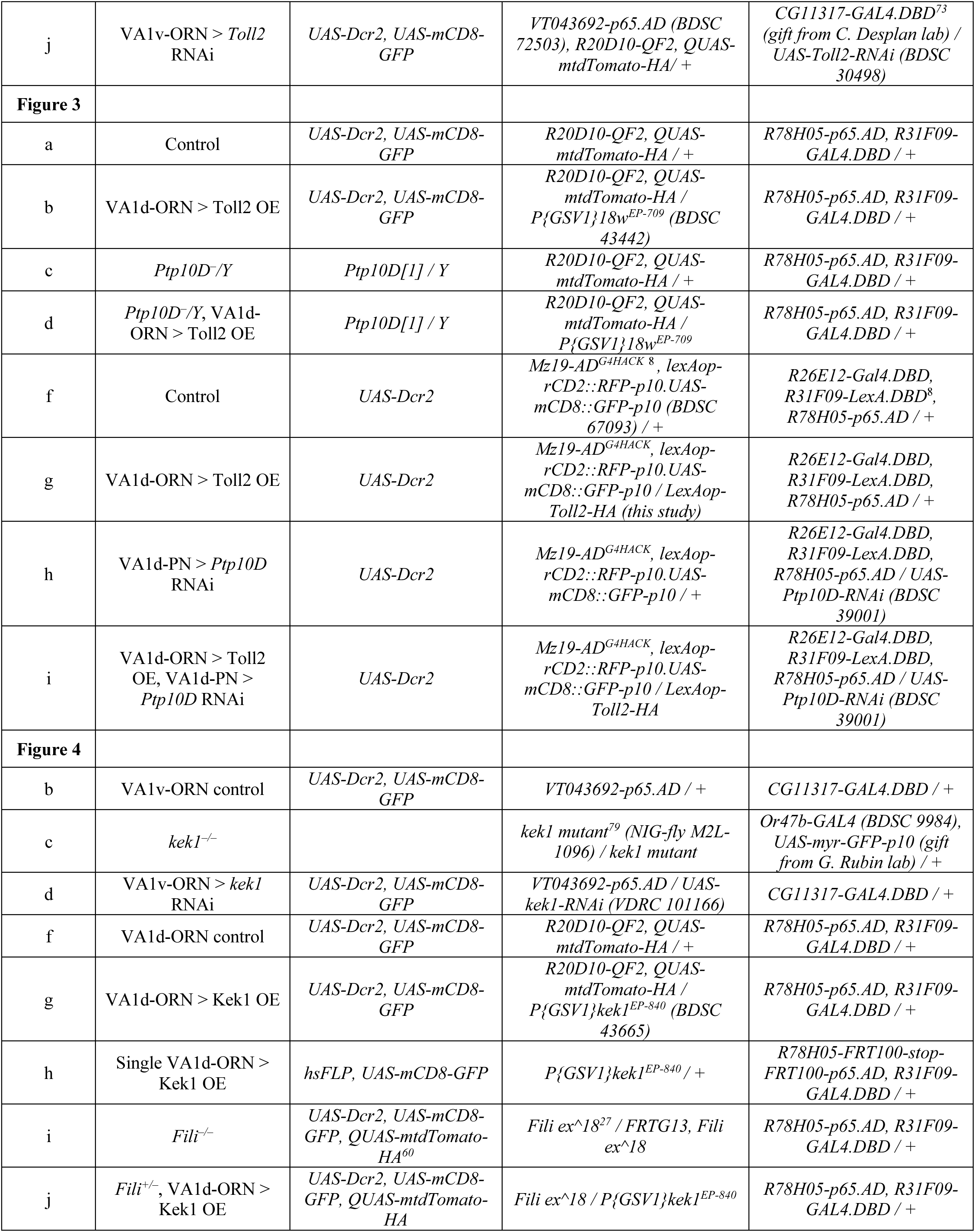
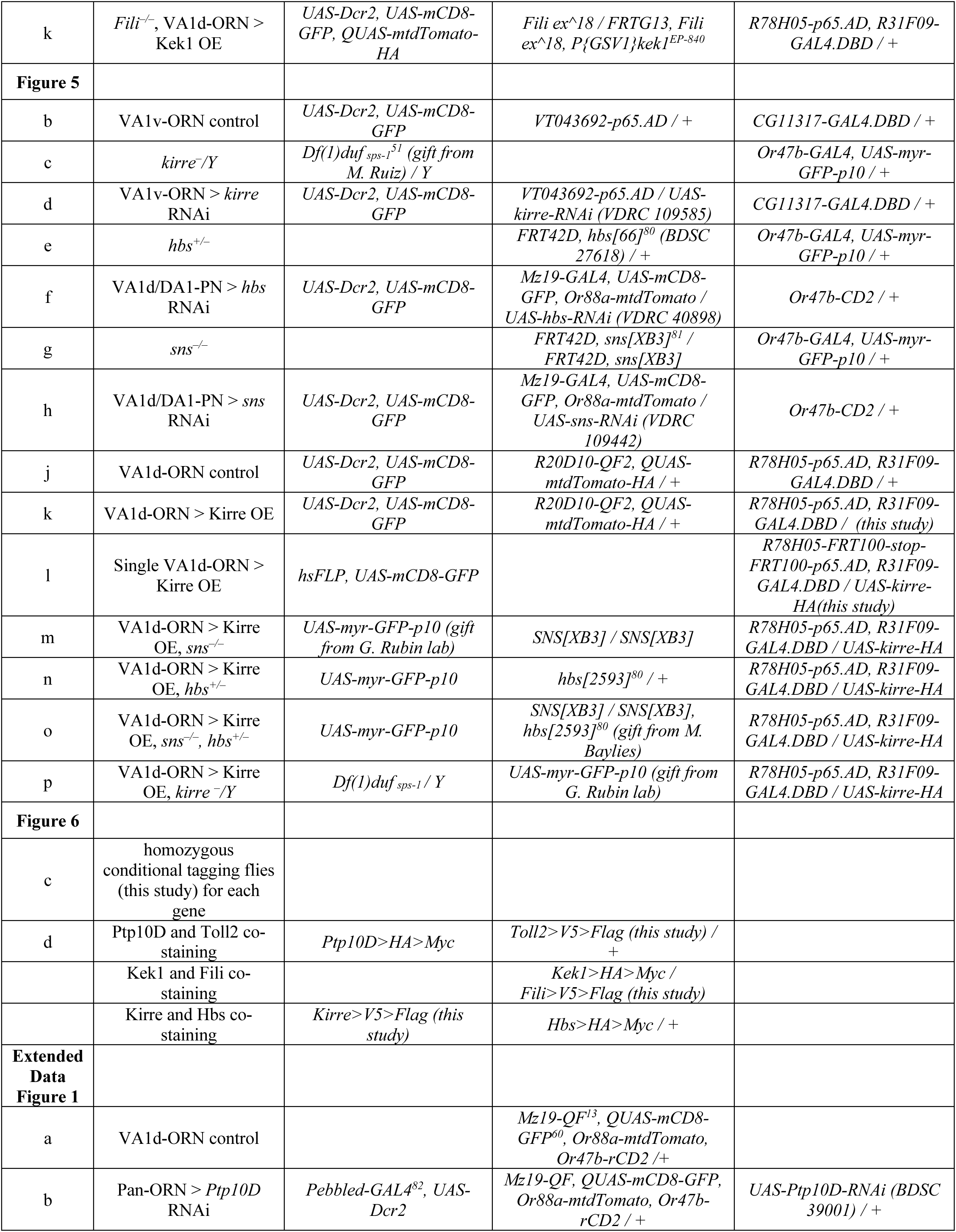
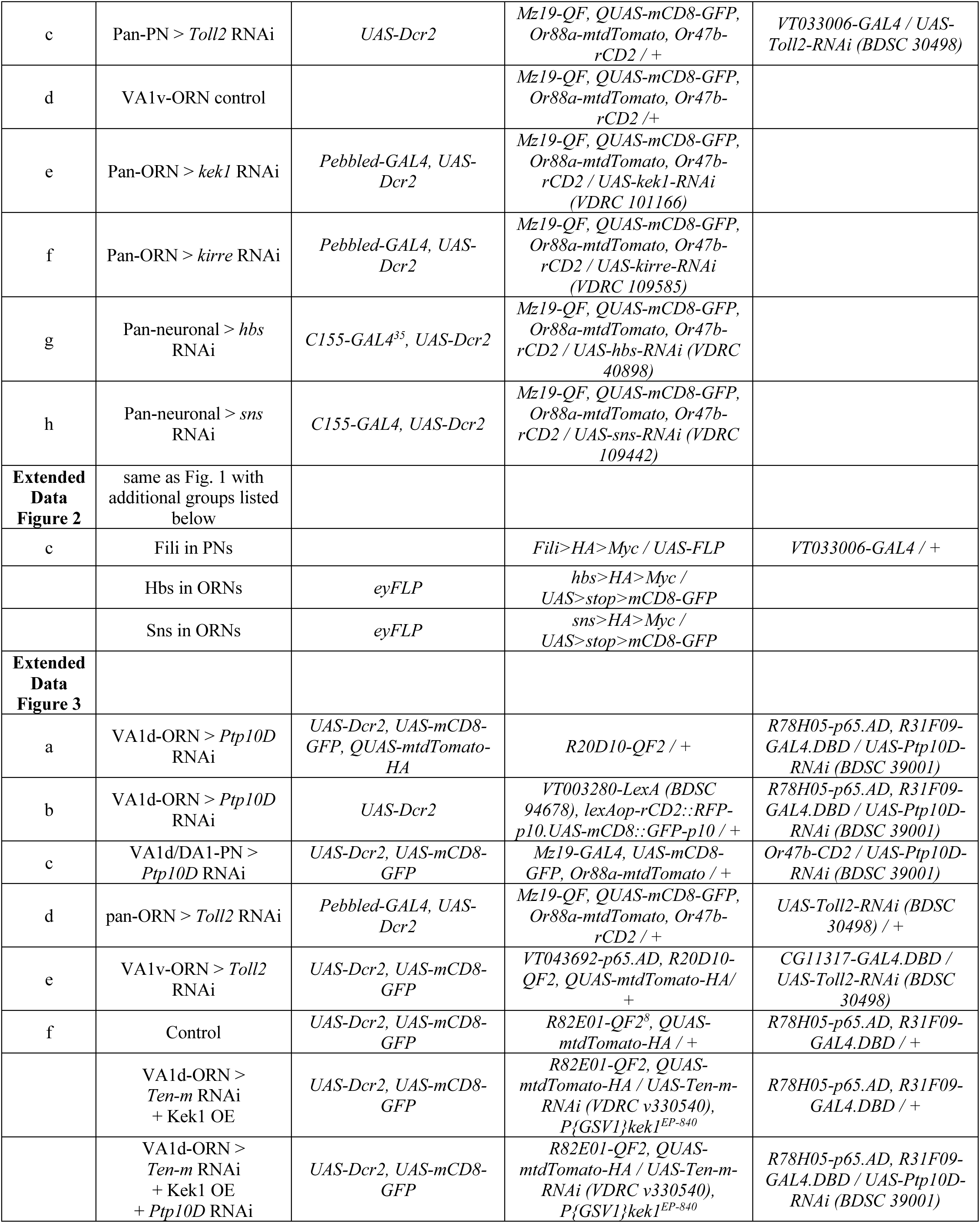
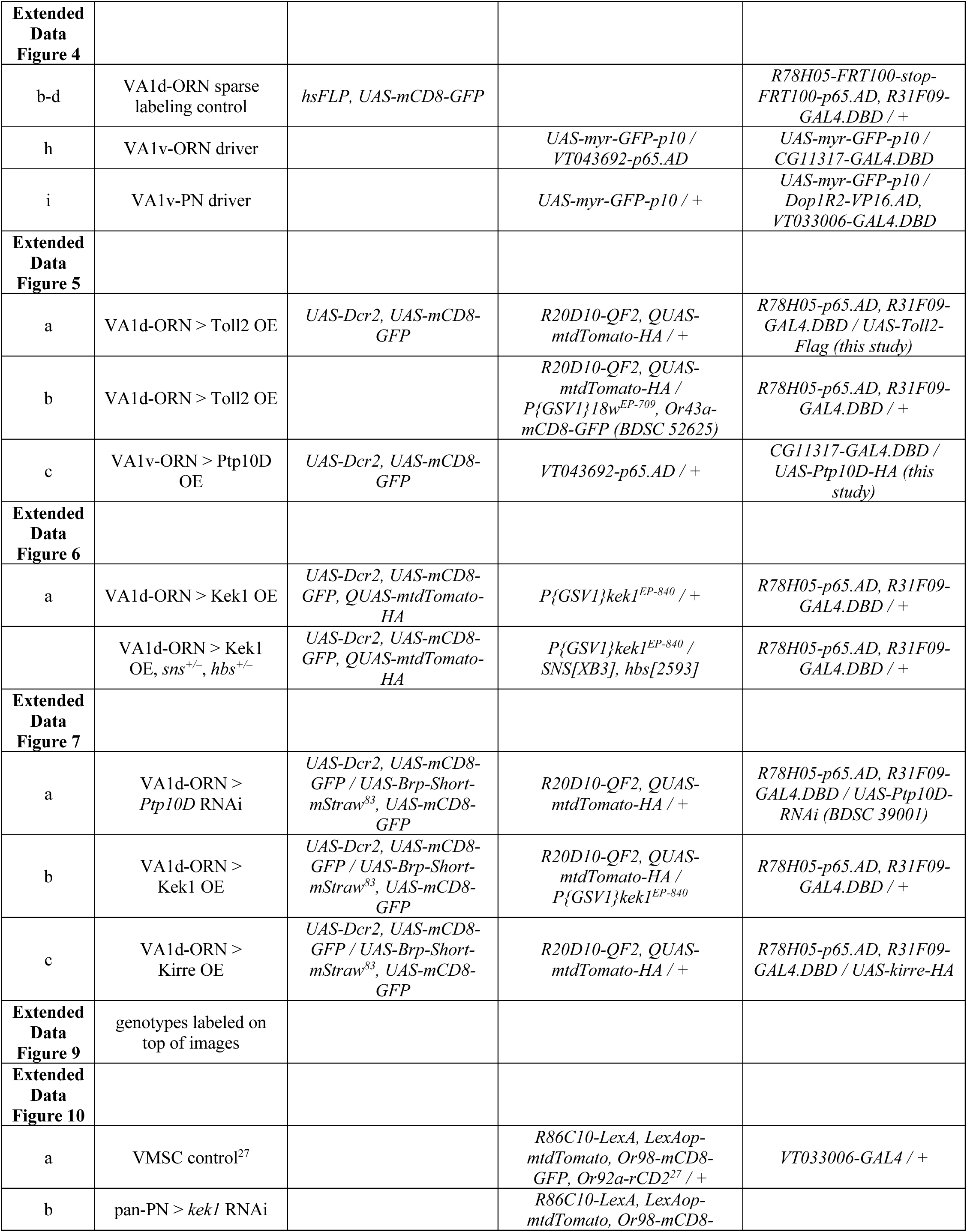
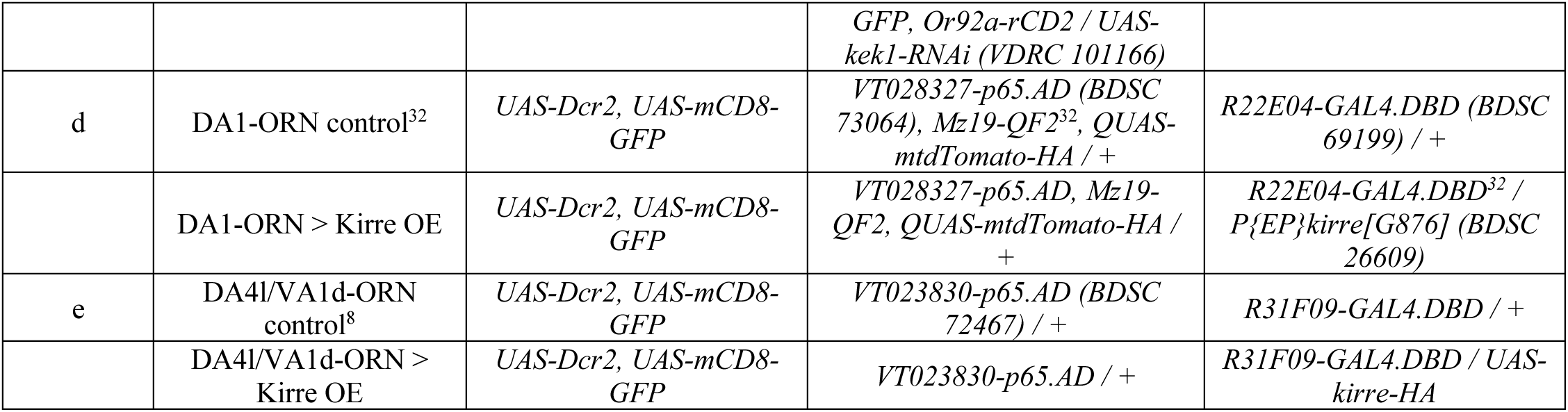
Summary of genotypes used in each experiment, arranged according to figure panels.

## Supplementary Materials

**Supplementary Videos 1–7.** Expression patterns of the 7 cell-surface proteins (CSPs) in the entire *Drosophila* pupal brains (42–48 hours after puparium formation) based on HA staining using newly generated knock-in fly strains. The name of the CSPs is labeled on the top right corner. Each video has two fly-through series from anterior to posterior along the *z*-axis (1-µm interval) of the confocal images. The first series show HA staining (cyan for Toll2, Fili, Hbs, and Sns; red for Ptp10D, Kek1, and Kirre) and N-cadherin staining (a neuropil marker) in blue. The second series is HA staining alone. The brightness of anterior, middle, and posterior sections was adjusted differently to achieve optimal visualization. Scale bars = 50 µm.

## Notes

### Competing Interest Statement

The authors have declared no competing interest.

### Summary of Updates

Organization of the paper and many more control experiments.

